# EEG Source Localization with High-Resolution 40-Tissue Head Models: A Comparative Study with MEG during Photic Driving

**DOI:** 10.64898/2026.11.20.768219

**Authors:** Guillermo Núñez Ponasso, Derek A. Drumm, Hannes Oppermann, Gregory M. Noetscher, Burkhard Maess, Thomas R. Knösche, Jens Haueisen, Sergey N. Makaroff

## Abstract

While MEG source localization pipelines often employ head models of one single compartment with homogeneous conductivity, researchers and clinicians use at least 3 different layers in head models for EEG source localization. Recent AI-based segmentation makes it possible to generate individualized head models with 40 high-resolution, non-nested tissues. We present a pipeline that can make use of, and test, these highly realistic models for EEG source localization: with the boundary element fast multipole method (BEM-FMM), we compute transcranial electric stimulation (TES) solutions on the full 40-tissue geometry and use them reciprocally to obtain the EEG leadfields. Our approach runs on a single modest workstation, requiring approximately 4.5 min per EEG channel for a mesh of about one million elements at a relative residual of 10^*−*4^. We applied the pipeline to six participants undergoing intermittent photic stimulation at half their individual alpha peak frequency, comparing 61-channel EEG source localizations from 40-tissue and 5-layer models with each other and with localizations from simultaneously recorded MEG. We further compared noiseless synthetic EEG topographies of the two models across the entire cortex. The two models produced highly consistent EEG source localizations (weighted relative difference below 3%), both in close agreement with the MEG localizations within the visual cortex. The synthetic topographies also agreed closely, with differences scaling approximately linearly with the angle between the source orientations of the two models, underscoring the importance of accurate source normals in fixed-orientation source models. These results establish 40-tissue models as a practical and robust basis for EEG source localization, and open the way to applications where nested geometries cannot represent the anatomy, such as infants or patients with skull defects, injuries, or tumors.

## 1 Introduction

Source localization, also known as source estimation or source reconstruction (Knösche and Haueisen 2022) consists of estimating the distribution of neural activity on the cortex that best explains a given set of neurophysiological recordings, either from electroencephalography (EEG), magnetoencephalography (MEG), or other modalities. EEG constitutes the most affordable and widely available modality of recordings; as such, source localization based on EEG is of great importance for both clinical and research-focused applications of neuroscience, including pre-surgical evaluation of patients with drug-resistant epilepsy (Brodbeck et al. 2011; Sharma et al. 2018) and brain-computer interfaces (Edelman et al. 2019).

While some information about the sources of EEG can be obtained from an analysis of the to-pography, which depends only on the measured signal, better estimates can be given by using volume conduction models solving the quasi-static Maxwell equations. This is the so-called *forward step* of source localization, where a relationship between sources and EEG topographies is established. The simplest of such forward models is the concentric spheres model (Papargiri et al. 2021), which approximates the major compartments of the human head (skin, skull, cerebrospinal fluid (CSF), and brain) using concentric spheres and has the advantage of having an analytic solution. To include individual head anatomy, researchers and practitioners generate head models from magnetic resonance imaging (MRI). Approaches like the overlapping spheres model (Huang et al. 1999) can incorporate a degree of anatomical information while still employing local analytical solutions. However, a forward model which makes full use of the head anatomy cannot be solved analytically and requires the computation of a numerical approximation to the solution.

Major source localization packages like the MNE Software (Gramfort et al. 2013; Gramfort et al. 2014), FieldTrip (Oostenveld et al. 2011), or Brainstorm (Tadel et al. 2011) implement either the boundary element method (BEM) or the finite element method (FEM) to obtain numerical solutions. In either case, these solvers are direct: meaning that the EEG/MEG sensor outputs are simulated from specified dipole locations and moments inside the head volume. Both BEM and FEM operate by constructing and subsequently inverting a so-called system matrix, which depends only on the anatomical models extracted from the MRI segmentation and on a choice of conductivity values for each tissue. Because the BEM system matrix is dense, this method is limited to low resolution head models. On the other hand FEM produces a sparse system matrix, and thus it can compute solutions involving more mesh elements. However, the FEM meshes are volumetric and a higher number of elements is needed to accurately describe the head geometry. Another disadvantage of FEM is that one cannot prescribe arbitrary incident fields, like those of point current dipoles, and needs instead to approximate them within the FEM mesh appealing to the Saint-Venant principle (Buchner et al. 1997).

The boundary element fast multipole method (BEM-FMM) (Makarov et al. 2018) combines the charge-based formulation of the BEM (Barnard et al. 1967; Núñez Ponasso 2024) with the fast multipole method (Greengard and Rokhlin 1987). This method has all of the advantages of BEM without the limitations of its dense system matrix, as the coupling with FMM renders it matrixless. This is both the main strength and weakness of the BEM-FMM, since, while the FMM acceleration allows us to carry out forward computations with models of a complexity previously unmatched — like those of neuromorphologically realistic microscopic neuronal arbors (Qi et al. 2025; Noetscher et al. 2026b) —once the incident field is changed the entire BEM-FMM iteration needs to be recomputed.

This limitation of BEM-FMM makes it unsuitable for the *direct approach* to forward modeling often used in BEM and FEM, where the EEG or MEG measurements are simulated for every dipole of the source model one at a time, as the number of dipoles in a typical source model ranges between 10,000 and 50,000. We overcame this limitation by coupling the BEM-FMM with electromagnetic reciprocity and applied it to MEG source localization with high-resolution 5-layer models (Núñez Ponasso et al. 2025; Drumm et al. 2026b). Electromagnetic reciprocity relates the solution of the EEG forward problem to the solution of a transcranial electric stimulation (TES) forward problem, and likewise it relates MEG to the solution of a transcranial magnetic stimulation (TMS) problem. This alternative approach, called the *reciprocal approach*, involves computing as many TES or TMS solutions as there are channels in the recording system. Since the number of channels typically ranges between 20–600, computing this number of stimulation solutions is feasible. The reciprocal approach has the added advantage that the source space can be put in one-to-one correspondence to elements of the BEM mesh, which in turn allows us to incorporate millions of sources into the source model, given that the BEM-FMM can feasibly carry out computations with millions of mesh elements.

To date, all major source localization software packages mentioned above employ 3–5 nested layer models for EEG source localization. The BEM with three layers and 10,000–20,000 sources based on the FreeSurfer (Fischl et al. 2004) automated segmentation is one of the most commonly used head models. The headreco 5-layer automated segmentation in the SimNIBS software package (Nielsen et al. 2018) has been used and tested against 3-layer models in our previous work (Núñez Ponasso et al. 2024; Núñez Ponasso et al. 2025; Drumm et al. 2026b) and similar 5-layer models have been employed in FEM source localization (Medani et al. 2023). Recently, AI-based automated segmentations of high-resolution non-nested 40-tissue head models have been introduced (ZMT Zurich MedTech AG / IT’IS Foundation, Zurich, Switzerland 2024). Non-nested head models can represent anatomical features that nested models cannot; this is relevant in EEG source localization for infants (Lew et al. 2013).

In this paper, we employ these head models in source localization for the first time and evaluate their agreement with nested 5-layer headreco models using both real and synthetic data. The aims of this paper are to introduce a TES methodology for reciprocal use in EEG source localization, to show the viability of source localization employing 40-tissue head models, and to compare these new head models to 5-layer head models for use in source localization. The main contributions in this paper are the following:

1. Introduction of a BEM-FMM pipeline for EEG source localization by coupling with electromagnetic reciprocity — previously, this was limited to MEG source localization.
2. Implementation of the Sim4Life 40-tissue models in both EEG and MEG source localization. Because of our reciprocal approach, this entailed the implementation of the same 40-tissue models for their use in TES and TMS.
3. Waveform and source analysis of joint EEG/MEG data from intermittent photic stimulation at ½ of the individual alpha peak frequency of a cohort of 6 healthy participants. The stimulation frequency was 5Hz on average, hence the evoked potentials and fields belong neither to the realm of visual evoked potentials (VEPs) nor steady-state visual evoked potentials (ssVEPs). Our analysis complements that of (Oppermann et al. 2025), which performed source analysis in the frequency space.
4. A comparison of noiseless synthetic EEG leadfields —and by reciprocity TES solutions — in the 40-tissue and 5-layer models.

## 2 Materials and Methods

### Participants and ethics

We made use of the dataset detailed in Oppermann et al. 2025 and Oppermann and Haueisen 2026, which consisted of a photic driving experiment using individualized intermittent photic stimulation (IPS) with stimulation frequencies set to half of the individual alpha peak frequency (iAPF), 4-6 Hz, of each participant. The stimulation was divided into 20 trials, with 80 flashes per trial (50% duty cycle) and a 5-second break between consecutive trials. Hence, a total of 1600 single stimuli were provided. Twelve healthy participants (5 female, 10 right-handed) with a mean age of 29±7 years participated in the study. The Ethics Committee of the Medical Faculty of Friedrich-Schiller University Jena approved the study, and all participants gave their written informed consent to the study and the use of the recorded data.

At the time of generation of the automated 40-tissue segmentation, 6 out of the 12 participants had to be removed because the resulting segmentation was faulty, with skin tissue being incorrectly labeled as muscle tissue in large portions of the skin surface, preventing us from assigning the correct tissue conductivities at the electrode locations in their corresponding models.

### Data Acquisition

In a first session, 5 minutes of resting-state EEG were recorded to determine the iAPF. Participants were seated in a relaxed position with their eyes closed. A 64-channel EEG cap (Ag/AgCl electrodes, waveguard™ original CA-212.s1, ANT Neuro b.v., Hengelo, Netherlands) and a mobile 64-channel referential DC-EEG amplifier (eego™ amplifier EE-225, ANT Neuro, b.v., Hengelo, Netherlands) were used for the recordings. A central electrode (5Z) served as an online reference, the electrical ground was placed on the left earlobe, and the sampling rate was set to 2048 Hz.

The second session comprised a combined EEG/MEG experiment with the individualized IPS. Participants were seated in a relaxed position with their eyes closed in a magnetically shielded room. EEG during stimulation was acquired employing a MEG-compatible cap comprising 61 Ag/AgCl electrodes (waveguard™ CA-169, ANT b.v., The Netherlands), placed according to the extended 10-20 system, with an online reference at the position AFz, the electrical ground on the left mastoid, and a sampling rate of 1000 Hz. The EEG amplifier provided by a MEG system (MEGIN, Helsinki, Finland) was used for the recordings. We used a Polhemus 3D digitizer to obtain the anatomical landmarks (nasion and both preauricular points), the individual locations of the EEG electrodes, and randomly distributed points over the head surface.

The data analyzed in this study pertain only to the second session.

### MRI

T1-weighted MRIs from all participants were acquired using a 3T Magnetom Prisma MRI scanner with a 0.8 mm resolution (Siemens Healthineers AG, Forchheim, Germany).

### Individual alpha peak frequency estimation

The resting-state EEG recording was only used for the determination of the iAPF. After bandpass filtering of the data (2–45 Hz, Butterworth, 4th order), we visually identified and removed any bad channels, before re-referencing the data to common average reference (CAR). The number of channels was reduced to eight occipital channels and the signal was divided into non-overlapping 30-second epochs. The first and last epochs were then removed. Subsequently, the power spectral density (PSD) was calculated using the Welch method (Welch 1967) for each occipital channel and segment. The Savitzky-Golay smoothing filter (Savitzky and Golay 1964) was applied and the iAPF was determined as the main peak in the alpha band (8-13 Hz) from the mean across the eight channels and segments.

### IPS data

We filtered the EEG recordings with a finite-impulse response (FIR) bandpass filter between 2 and 45 Hz for visual inspection and removal of bad channels and bad trials, i.e. those with flat lines or high amplitudes, and channels with repeating artifacts. Subsequently, we applied independent component analysis (ICA) to detect and reduce the impact of electrocardiogram (ECG) and eye-movement artifacts. The cleaned EEG data were re-referenced to CAR. For further source localization tasks, the recordings were divided into epochs. First, stimulation epochs were created, containing 250 ms of signal, starting 50 ms prior to the onset of each stimulation. Depending on the number of trials removed, 13 – 20 epochs were averaged and used for subsequent source reconstruction. Second, baseline (non-stimulation, resting state) epochs were created. Epochs were taken from the breaks between each trial and lasted 3.5 seconds. A maximum of 20 epochs was extracted from the data.

### Signal peak detection

Filtered and averaged EEG signals from intermittent photic stimulation were inspected manually and observed to be periodic with a period corresponding to the stimulation frequency (one-half of the iAPF). Four distinctly visible peaks, two positive and two negative were identified in all participants from a visual inspection of the midline, occipital Oz channel signal. We define N_II_ as the negative peak of largest magnitude. There is a consistent pattern across signals where N_II_ is immediately followed by a high magnitude positive peak; we label this as P_I_. We defined the beginning of the signal period at P_I_, shifting the signal by the periodicity if necessary. With this convention, the succession of peaks is P_I_, N_I_, P_II_, and N_II_, with N_I_ and P_II_ being the peaks of negative and positive polarity occurring between P_I_ and N_II_, respectively. For comparison among participants, we remapped every EEG signal onto a [0, 1] phase space so that the P_I_ peak occurs at *t* = 0 (and *t* = 1, since the waveform is periodic), we refer to this as normalized phase space (NPS). We then compute the participant-averaged signals within the NPS. We perform this same procedure for the MEG signals, mapping to NPS using the latency of the EEG P_I_ peak. The signal strengths are then normalized by the strength of the signal at the P_I_ peak.

### Head modeling: 40-tissue models

Surface meshes of 40 tissues with non-nested topology were extracted using the Sim4Life AI-based anatomy modeling pipeline *head40* (ZMT Zurich MedTech AG / IT’IS Foundation, Zurich, Switzerland 2024) and assigned material properties from the low-frequency conductivity section of the IT’IS Tissue Properties Database (Baumgartner et al. 2025). We used the algorithm for surface extraction provided by *head40* which ensures that every interface between adjacent tissue compartments, except for those at the domain boundary, is represented by a pair of coincident but inversely oriented surface faces, with one face belonging to each adjacent compartment. Duplicate faces were removed and the conductivity contrast associated with each remaining facet is then assigned according to the orientation of its outward normal vector, with the corresponding inner and outer conductivities defined consistently with respect to that orientation. The resulting models contain approximately 1 million facets in total with an average edge length of 1.5 mm.

### Head modeling: 5-layer models

MRI T1-weighted data for the 6 healthy participants were automatically segmented using the *headreco* pipeline (Nielsen et al. 2018) of the SimNIBS software package (Saturnino et al. 2019). This pipeline provides five major nested compartments: skin, skull, CSF, gray matter (GM), and white matter (WM), as well as two smaller compartments for the eyes and the ventricles. Because the resulting BEM meshes are nested, all triangles were oriented along the outward normal, and each triangle was assigned the conductivities inside and outside the tissue whose boundary it belongs to. All 5-layer models were refined by a global barycentric subdivision of all triangles, which resulted in approximately 800,000 triangles per model. The tissue conductivities were set to the values used in the 40-tissue models, with one exception. In the 5-layer models, the skin surface encloses a volume that the 40-tissue models resolve into skin, muscle, fat, and other soft tissues. To better match the 40-tissue models, the skin compartment was therefore assigned the mean conductivity of these enclosed tissues. This average was computed as the unweighted mean of the conductivities of skin, muscle, and the remaining soft tissues of the 40-tissue model, (0.148 + 0.078 + 0.461)*/*3 *≈* 0.229 S m^*−*1^.

### Further post-processing for both head models

Both the 40-tissue and 5-layer models were smoothed using a non-manifold global Taubin smoothing (Taubin 1995) with positive scale factor *λ* = 0.5 and negative scale factor *µ* = *−*0.53.

### Source space

The source space for both the 40-tissue and 5-layer models consists of point current dipoles located on the triangle centers of all triangles in the gray-matter/white-matter interface. The orientation of every dipole is given by the outward-pointing unit normal to the triangle facet where each dipole sits. The dipole moment for each triangle is set to 1mA m. The number of dipoles in the source model of each participant in the 40-tissue head models is of approximately 125,000*±*10,000, and in the 5-layer model is approximately 220,000*±*25,000.

### Forward solver: BEM-FMM

All forward numerical calculations were performed using the boundary element fast multipole method (BEM-FMM) (Makarov et al. 2018), which consists of the application of the fast multipole method (FMM) (Greengard and Rokhlin 1987) to the charge-based formulation of the BEM (Gelernter and Swihart 1964; Barnard et al. 1967). Given a conducting volume with piece-wise constant isotropic conductivity *σ* in every compartment and an impressed electric field **E**^*i*^, the integral equation for the charge density *ρ* on the surfaces of discontinuity *S* of the conductivity is the following:

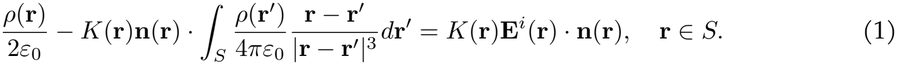

Here, *K*(**r**) = (*σ−* (**r**) *− σ*_+_(**r**))*/*(*σ−* (**r**) + *σ*_+_(**r**)) is the conductivity contrast at the surface point **r**. This is a Fredholm integral equation of the second kind. After discretization of the surfaces *S* by a triangular mesh, we apply the Galerkin method with zeroth-order (or pulse) basis functions to convert the integral equation into the linear equation

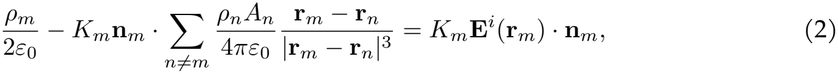

where **r**_*m*_ and *A*_*m*_ are the center and area of triangle *t*_*m*_, respectively. The summation term is accelerated with the FMM implemented in the FMM3D library of The Flatiron Institute 2012. The expression in (2) is used to formulate an iterative generalized minimal residual method (GMRES) for the calculation of the discretized charge density *ρ*_*m*_; see (Núñez Ponasso 2024). This iteration is terminated when the GMRES achieves a relative residual within a certain threshold. In this study, the GMRES threshold was taken to be 1 *×* 10^*−*4^.

### Forward solution: reciprocal approach

To calculate the forward EEG solution for our chosen source model, we apply Helmholtz reciprocity (Helmholtz 1853), which is a classical result that can be applied to relate the fields generated by a current dipole with the fields generated by an electrode pair.

Consider a current point-dipole at position **p** and moment **q** = *q***d**, where *q* = 1 [A] is the magnitude of the dipole current and **d** is the direction component of the moment, measured in [m]. Denote by *ϕ*_1_ the electric potential on the skin caused by the current dipole. Consider two current electrodes *e*_*i*_ and *e*_0_ centered at skin positions **r**_*i*_ and **r**_0_, respectively. Inject a total current of 1 [A] flowing into the skin from electrode *e*_*i*_ and a current of 1 [A] flowing out of the skin from electrode *e*_0_. Denote by **E**_2_(**r**; *e*_*i*_) the electric field elicited by this TES arrangement with electrode pair (*e*_*i*_, *e*_0_) on the cortical location **r**. Then, Helmholtz Reciprocity states (cf. Plonsey 1963):

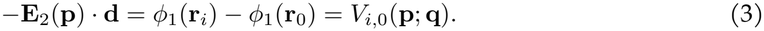

For a derivation of this reciprocity relation based on the quasi-static approximation to Maxwell’s equations, see Supplement 1. The equation above is a simplification for point electrodes. While the point electrode model is sometimes used in practice (Knösche and Haueisen 2022), we note that a version of this equation for general contact-impedance electrodes can be derived similarly (although with more technical details). Supplement 1 also contains an alternative derivation based on circuital reciprocity.

### TES modeling approach

To model TES using the BEM-FMM, we used the approach detailed in the supplementary material of Makarov et al. 2021 and Drumm et al. 2026a, where the incident field **E**^*i*^ in Equation (1) is set to zero and boundary conditions of either Dirichlet, Neumann, or Robin type are set. In this work, we used voltage electrodes which are specified by an additional Dirichlet boundary condition and a pre-conditioner is employed at the electrode locations in order to solve the linear system — also built using FMM acceleration. The use of zeroth-order basis functions in the BEM causes approximation errors which need to be corrected. We achieve this by using a near field correction employing a large number (*K* = 1024) of neighbor integral corrections for every triangle center. Namely, each term in the sum

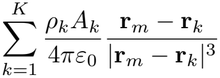

which involves the *K* nearest neighbors to triangle *t*_*m*_, and contributes to the iterative calculation of *ρ*_*m*_, is substituted by the near field correction

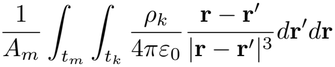

which is computed semi-analytically using the formulas of (Wilton et al. 1984) for the inner integral and Gaussian cubature for the outer integral. This calculation was implemented in the `C` programming language and wrapped into our `MATLAB` pipeline for increased performance. More on near field corrections in BEM-FMM can be found in Section 5 of (Núñez Ponasso 2024). We validated our near field correction approach (using 1024 neighbor integrals) against the adaptive mesh refinement (AMR) algorithm proposed in Drumm et al. 2026a, finding a relative difference of 1%. This AMR method has been validated against Sim4Life hexahedral and FEM solvers for deep (hippocampal) brain targets in Noetscher et al. 2026a, showing a relative difference of 1.7% and 1.2%, respectively.

### Source localization: Calculation of the leadfield matrix

The main ingredient of the inverse operator is the leadfield matrix, which for EEG source localization (Knösche and Haueisen 2022) using an EEG montage with *n* non-reference channels and *N* sources is the *n × N* matrix *L* whose (*i, j*)-th entry is given by

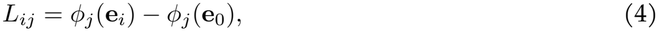

where *ϕ*_*j*_ is the scalp voltage induced by dipole *j*, **e**_*i*_ is the center of the *i*-th electrode (*i* = 1, …, *n*) and *i* = 0 corresponds to the reference electrode. By Equation (3) we have that

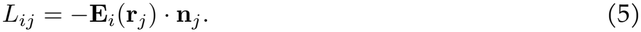

Due to the fact that the surface charge density is equal to the jump of the normal volume field in the surface (cf. section 2.3.5 of Griffiths 1999), and the incident field is zero, we approximate the leadfield matrix as

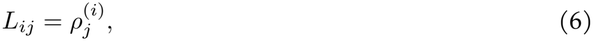

where 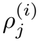 is the surface charge density at triangle *t*_*j*_ in the TES solution for voltage-driven electrodes *e*_*i*_ and *e*_0_ with voltages +1 V and *−*1 V respectively. The boundary condition for these voltage-driven electrodes is that the integral (or sum in the discretized case) of charge density the contact surfaces of *e*_*i*_ and *e*_0_ must equal +1 and *−*1 respectively. Note that, since a global scaling factor in the LFM does not affect the distribution of the inverse solution, the choice of unit and sign does not matter.

The EEG positions were manually co-registered to each participant’s head model using as guide-lines the left and right preauricular points, nasion, and the Polhemus digitization points. We then computationally imprinted the electrodes: we refined the triangles within a radius of 5mm of the co-registered electrode centers in order to approximate a circular contact region for the electrodes.

### Source localization: the inverse solution

The combination of Equations 3, 4, and 6 describe the linear relationship between surface charge density and the measured EEG signals, given as the forward model *Lx* = *y*, where *x* is the *N × T* array of neural dipole source strengths and *y* is the *n × T* array of *T* signal measurements. This system is often solved by constructing an inverse operator using minimum norm estimation (Knösche and Haueisen 2022; Drumm et al. 2026b):

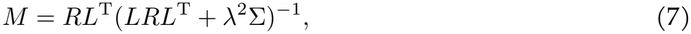

where *R* is the source covariance prior, Σ is the noise covariance of the measured signals, and *λ* is a regularization parameter, typically related to the signal-to-noise ratio. The inverse solution can then be recovered as *x* = *My*. However, there are two forms of bias that must be corrected in this solution. First, the presence of a common reference electrode across all rows of *L* as seen in Equation (4) leads to a common mode across all basis functions and thus a bias in the localized sources. This bias is most easily removed by removing the average basis function **m** (that is, the average of the rows of *L*) from the leadfield, 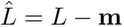. Second, the solution is biased due to the presence of signal measurement noise. To alleviate this bias, we instead solve the whitened linear system 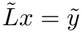, with 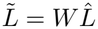 and 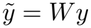, where *W* = Σ^*−*1*/*2^ is a whitening operator. By applying *W* to the linear system, the measured signals are mapped so that the new whitened signals all have covariance 1. In this manner, the whitened inverse operator is

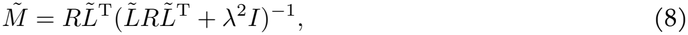

and the inverse solution is recovered as 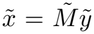. We used Equation (8) to compute the localized sources from the identified EEG peaks across all participant models.

Because the distance between the EEG electrodes and the white matter is smallest on the gyri, the source localization solutions will show strong activation on more superficial sources. We bias our solutions away from this superficial activation through the prior *R* on the source space, taken as a diagonal matrix whose entries are the column norms of the leadfield: 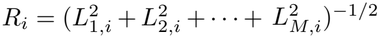. We further bias our solutions away from superficial regions onto the most statistically significant dipoles using dynamic statistical parametric mapping (dSPM) (Dale et al. 2000), which normalizes the localized sources by:

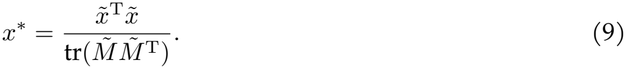

Finally, we rescale all normalized solutions to lie within [0, 1]. All source localization solutions shown in this work may be assumed to be dSPM normalized and rescaled, unless directly stated otherwise.

### Source localization metrics

We compute the MEG leadfields using charge-based BEM-FMM (Núñez Ponasso et al. 2025; Drumm et al. 2026b) and compute the inverse operator in the same manner as the EEG leadfield using Equation (8). Denote by *X* and *X*^*′*^ the EEG and MEG source localization solutions, respectively. We compute the weighted centroid of activation for each solution by

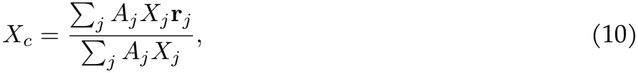

where *j* is taken over all triangles such that *X*_*j*_ *> t* for some activation threshold *t* (we will use an activation threshold of *t* = 0.8 for all solutions in this work unless stated otherwise). We then consider two difference metrics for comparing the EEG and MEG localizations. First, we use the distance between the EEG and MEG centroids of activation, 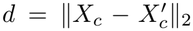. This distance is computed per hemisphere, except when the activation threshold is not met on a given hemisphere for *X* or *X*^*′*^; in that case, the comparison is removed. Second, we used an area weighted relative difference percentage,

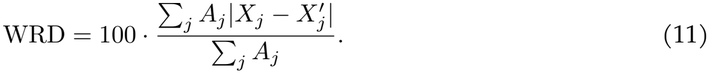

Because our MEG methodology is well tested, we will use these metrics to determine the validity of our EEG localizations.

Finally, we also compare the differences in localized EEG sources between the 40-tissue and 5-layer models using the same metrics as in Equations 10 and 11. For visualization, we will plot the solution centroids in the participant-averaged MNI coordinate space (Evans et al. 1993). Cross-comparison between these models on participant-specific coordinates requires additional co-registration, as detailed below.

### 40-tissue and 5-layer cross-model comparison

The major difference between the 40-tissue and 5-layer model source spaces is that the GM-WM interface of the 5-layer model includes the cere-bellum. To adequately compare both source spaces, we first removed cerebellar sources from the 5-layer model by co-registering both source spaces with the iterative closest point (ICP) algorithm (Bergström and Edlund 2014) and then excluding source locations far from all sources of the 40-tissue model. The distance threshold used to remove cerebellum points was 6mm across all participants. A second ICP step was then performed to further align both source spaces.

We compared the differences in synthetic noiseless topographies of the 40-tissue and 5-layer models for all dipoles in each source model using the described 61-channel waveguard CA-169 EEG cap. The topography **T**_*j*_ for dipole *j* is defined as the *j*-th column of the leadfield matrix, i.e. 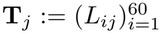, where *L* is calculated according to Equation (6). In what follows, we consider the source space of the 40-tissue model as a reference for comparison.

Since both source models cannot be perfectly co-registered, for every dipole *p*_*j*_ in the reference source space we selected *k* = 6 nearest dipoles in the 5-layer model among those with an angle of at most *π/*8 radians with respect to *p*_*j*_. The topographies of these *k* dipoles were averaged and the discrepancy of this averaged topography with respect to the topography of *p*_*j*_ was calculated. The measures of discrepancy we used were RDM and lnMAG (Meijs et al. 1989), defined for topographies **T**_1_ and **T**_2_ as follows

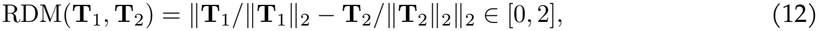

and

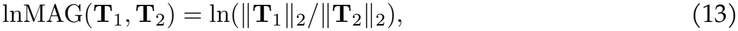

where ∥**x** ∥_2_ is the Euclidean norm of **x** ∈ R^*n*^. To remove the effect of global scale differences in the estimation of the magnitude variation, we consider the following:

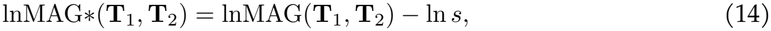

where *s* is the global scale factor, i.e. the median of the set of quotients (**T**_1_)_*i*_*/*(**T**_2_)_*i*_, for all *i* = 1, …, *n* whenever (**T**_2_)_*i*_ ≠ 0. For every topography **T**_*j*_ *∈* R^60^ of dipole *p*_*j*_ in the source space of the 40-tissue model, we calculated 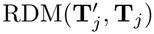 and 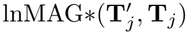, where 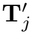 is the averaged topography near dipole *p*_*j*_ in the 5-layer model. Figure 1 illustrates the topographical comparison approach.

**Figure 1:**
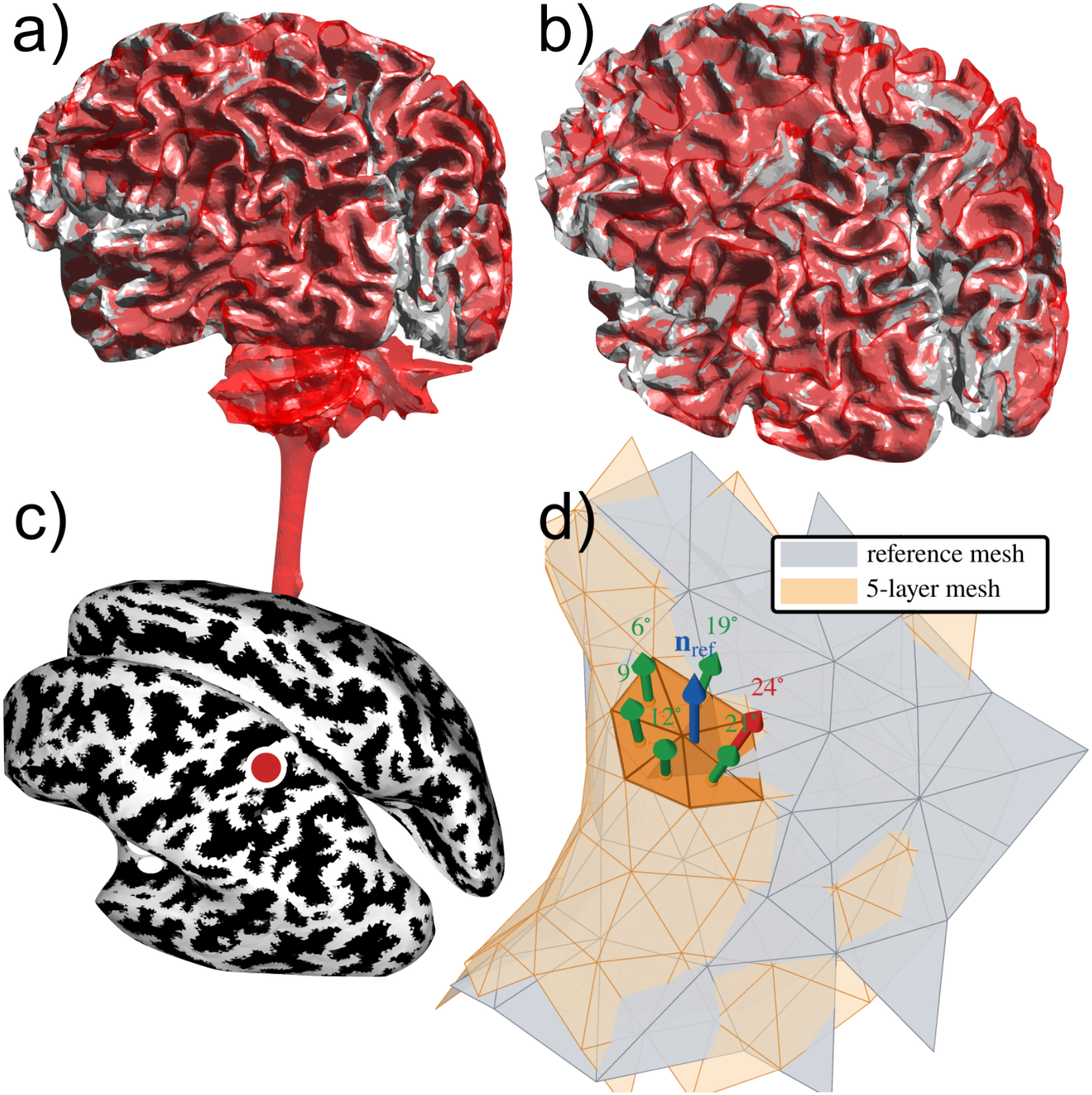
a) Initial co-registration of both source spaces, the 5-layer headreco mesh with cerebellum and brain stem is displayed in red; b) co-registration after truncating the cerebellum from the 5-layer source space and performing ICP; c) inflated source space with curvature map, white regions correspond to gyri and black regions to sulci. The location within the brain of the reference source in d) is indicated with a red circle; this corresponds to a gyral point in the postcentral gyrus; d) close-up image of a reference source in the 40-tissue model with near triangles in the 5-layer mesh and corresponding normal. In blue, we indicate the direction of the reference source, in green we indicate the directions of those near-sources within an angle of *π/*8 radians (approx. 23°) of the reference source (used in the average), in red we indicate the sources rejected due to having an angle over the threshold of *π/*8 radians.

To better assess which portion of the calculated differences is due to differences in the location and orientation of *p*_*j*_ and the average of its nearby dipoles, we computed a baseline RDM and lnMAG of the 40-tissue model against itself by taking the average topography 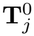 of the *k* = 6 nearest sources to *p*_*j*_ in the 40-tissue model’s source space. We report the *excess RDM* (XRDM) given as

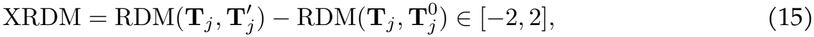

for every source in the 40-tissue model, as a measure of the signed discrepancy in topographies due to model differences. A higher magnitude of XRDM corresponds to larger topography differences.

### Computational resources for runtime statistics

We computed runtime statistics for the calculation of a TES solution on our `ECE-93` workstation: a Dell Precision 5860 Tower with an Intel(R) Xeon(R) w5-2445, 2096MHz processor with 10 cores and 20 logical processors, with 128GB of installed RAM, operating on Microsoft Windows 11 Enterprise. Pricing in November 2026 for this workstation was at $5,329.00.

## 3 Results

### Computational cost of the BEM-FMM forward modeling

Both model types (40-tissue, ~1 million *±*30, 000 elements; 5-layer, 820,000 *±* 70,000 triangles) had similar computational costs on our `ECE-93` machine, so the results are pooled. The runtime for each basis function (i.e. every channel pair), to a relative residual of 1 *×* 10^*−*4^, was 266.5 *±* 5.8s, corresponding to a processing speed of approximately 3,400 mesh elements per second (3,100–3,700 across the two model types). The precomputation of the near-field integral correction took 427 *±* 34s. The total time required to compute a 61-channel EEG leadfield matrix was therefore approximately 4 hours and 30 minutes. Storing the sparse near-field correction matrices required 58–65GB depending on the model, with about 1GB of additional memory needed for the rest of the execution.

### Waveform in the iAPF stimulation

Table 1 shows the determined peaks for the occipital Oz EEG channel, as well as the corresponding cycle and *α* stimulation frequencies, for all participants. Figure 2a shows the same occipital EEG channel signal, as well as the signal from left and right hemisphere magnetometer channels, mapped to NPS and averaged across participants. Figure 2b shows the participant-averaged EEG signals for all channels, plotted in NPS, with the Oz channel highlighted. Figure 2c shows the EEG topography of the participant-averaged signals at each of the four distinct peaks.

**Table 1:** The waveform frequency *f* of the occipital Oz channel is shown for each participant, with the corresponding *α* stimulation frequency for comparison (Oppermann and Haueisen 2026). Signal peak latencies and amplitude are also shown. Latencies are measured relative to P_I_ (so the P_I_ latency is 0 and *f* Hz for all participants and therefore omitted). The participant-averaged values and standard deviations are shown on the bottom row.

|  | Frequency [Hz] |  | P<sub>I</sub> | N<sub>I</sub> |  | P<sub>II</sub> |  | N<sub>II</sub> |  |
|---|---|---|---|---|---|---|---|---|---|
| Participant | *f* | *α*<sub>SF</sub> | *A* [µV] | *t* [ms] | *A* [µV] | *t* [ms] | *A* [µV] | *t* [ms] | *A* [µV] |
| P03 | 5.92 | 4.75 | 4.42 | 46 | −0.86 | 66 | 0.49 | 99 | −3.78 |
| P06 | 5.21 | 5.50 | 3.46 | 45 | −3.23 | 105 | 3.52 | 141 | −3.41 |
| P08 | 5.52 | 5.00 | 2.61 | 38 | −1.71 | 108 | 1.20 | 137 | −3.13 |
| P10 | 5.10 | 5.70 | 4.68 | 40 | −3.72 | 79 | 4.71 | 129 | −6.38 |
| P14 | 4.41 | 4.95 | 6.74 | 42 | −1.66 | 109 | 1.57 | 161 | −8.32 |
| P15 | 4.61 | 5.45 | 7.43 | 73 | −4.01 | 105 | 2.79 | 165 | −6.28 |
| Mean ± SD | 5.13 ± 0.56 | 5.23 ± 0.38 | 4.89 ± 1.86 | 47 ± 13 | −2.53 ± 1.29 | 95 ± 18 | 2.38 ± 1.58 | 139 ± 24 | −5.22 ± 2.09 |

**Figure 2:**
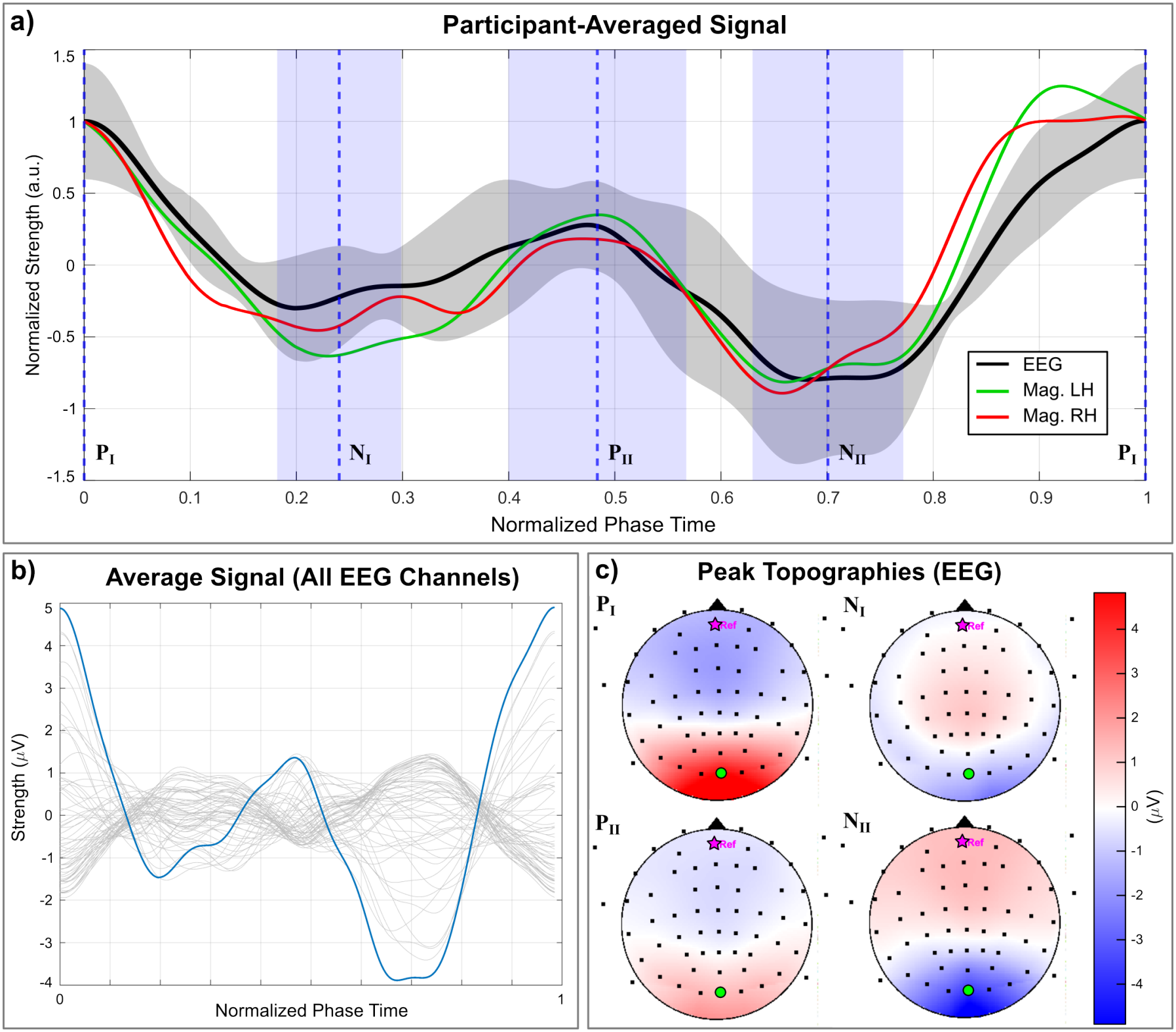
a) We show the Oz channel signal (black), as well as MEG signals averaged from a cluster of magnetometers on the left (green) and right (red) hemispheres, each averaged across all participants and plotted in normalized phase space. (Selected magnetometer channels and signals may be found in Supplement 2.) Peaks P_I_, N_I_, P_II_, and N_II_ are averaged across participants and plotted as blue dashed lines, with corresponding 95% confidence intervals. Shadowed regions indicate the 95% confidence interval of the participant-averaged EEG signal. b) The participant-averaged EEG signal is computed across all channels and plotted in phase space, with the signal from the Oz channel highlighted in blue. c) The averaged signal topographies are plotted at the P_I_, N_I_, P_II_, and N_II_ peaks. The anterior side of the head relative to the 61-channel electrode montage is indicated with a black triangle. The reference electrode is highlighted in magenta, and the Oz channel is highlighted in green.

### Source localization results: cross comparisons

Figure 4a shows example EEG and MEG source localization solutions and corresponding RDM percentage for participant P08 at the P_I_ peak. Figure 4b shows the per-hemisphere solution centroid distances between the EEG and MEG solutions across all participants and signal peaks, and Figure 4c shows the corresponding WRD percentages. Table 2 shows the EEG source localization comparison metrics between the 40-tissue and 5-layer models for each participant and signal peak. Figure 5 shows the EEG source localization solution centroids for the 40-tissue and 5-layer models visualized in the participant-averaged MNI coordinate space. Additional data tables corresponding to these results can be found in Supplement 3.

**Table 2:** Comparison of the 40-tissue and 5-layer head models for the EEG source localization solutions at each signal peak, with an activation threshold *t* = 0.8. For each participant, the centroid distance on the left (LH) and right (RH) hemispheres and WRD percentage between the two models’ solutions are shown. A dash (–) indicates that the activation threshold was not met for at least one of the models in that hemisphere. The participant-averaged values and standard deviations are shown on the bottom row.

|  | P<sub>I</sub> |  |  | N<sub>I</sub> |  |  | P<sub>II</sub> |  |  | N<sub>II</sub> |  |  |
|---|---|---|---|---|---|---|---|---|---|---|---|---|
| Participant | LH [mm] | RH [mm] | WRD [%] | LH [mm] | RH [mm] | WRD [%] | LH [mm] | RH [mm] | WRD [%] | LH [mm] | RH [mm] | WRD [%] |
| P03 | – | 1.0 | 1.71 | – | 24.8 | 2.38 | 12.5 | – | 2.14 | 9.3 | 11.4 | 2.88 |
| P06 | 20.4 | 2.1 | 2.88 | 12.0 | 1.0 | 3.04 | 4.0 | 6.0 | 3.04 | 0.5 | 0.7 | 2.96 |
| P08 | 10.9 | 19.7 | 2.43 | 2.9 | 2.8 | 2.30 | 8.1 | 5.6 | 3.12 | 4.4 | 18.0 | 2.31 |
| P10 | 19.6 | 49.5 | 5.59 | 20.5 | 45.3 | 4.98 | 10.0 | 2.6 | 4.70 | 7.2 | 6.7 | 4.49 |
| P14 | 3.8 | 16.0 | 1.88 | 25.1 | 11.0 | 2.58 | 8.7 | 11.9 | 2.86 | 6.2 | 14.5 | 1.78 |
| P15 | 21.4 | 6.6 | 2.37 | 34.0 | 15.6 | 2.27 | 22.7 | 14.2 | 2.07 | 0.6 | 1.7 | 2.05 |
| Mean ± SD | 15.2 ± 7.6 | 15.8 ± 18.1 | 2.81 ± 1.43 | 18.9 ± 11.9 | 16.7 ± 16.5 | 2.93 ± 1.05 | 11.0 ± 6.3 | 8.1 ± 4.8 | 2.99 ± 0.95 | 4.7 ± 3.6 | 8.8 ± 7.0 | 2.74 ± 0.97 |

**Figure 3:**
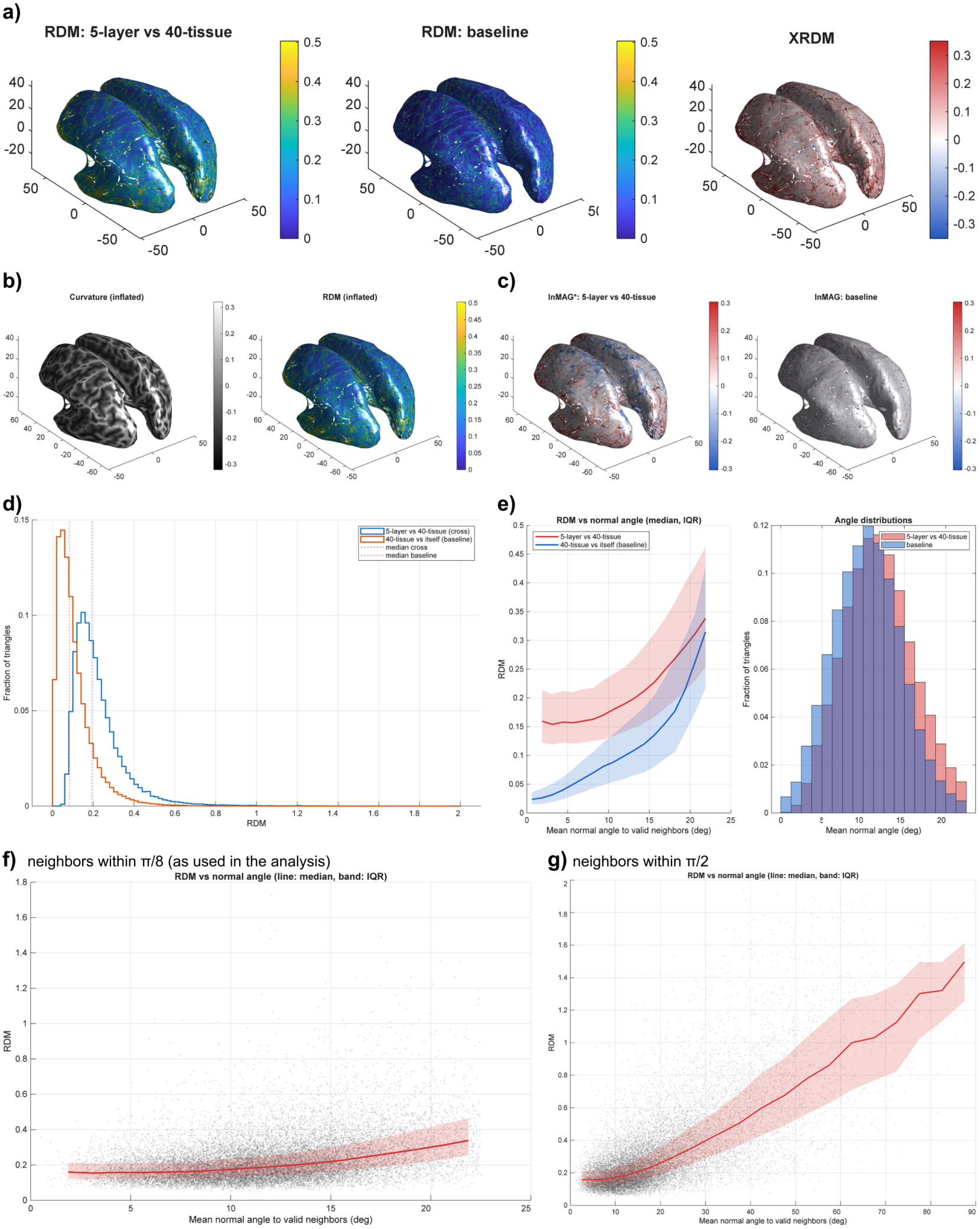
a) RDM in the cross (5-layer vs 40-tissue) comparison, together with the baseline RDM and excess RDM in the inflated white matter mesh of P03; b) RDM compared to the curvature map in the inflated mesh, light regions in the curvature map correspond to gyri, while dark regions correspond to sulci; c) lnMAG* in the cross-com2p0arison vs lnMAG in the baseline; d) a histogram of RDM in all analyzed reference triangles in the cross-comparison and baseline; e) RDM vs normal angles in the cross-comparison and baseline, toghether with a histogram of the mean normal angle; f) RDM against normal angle to centroid, plotted from 0 to *π/*2; g) RDM against normal angle to centroid; plotted from 0 to *π/*8.

**Figure 4:**
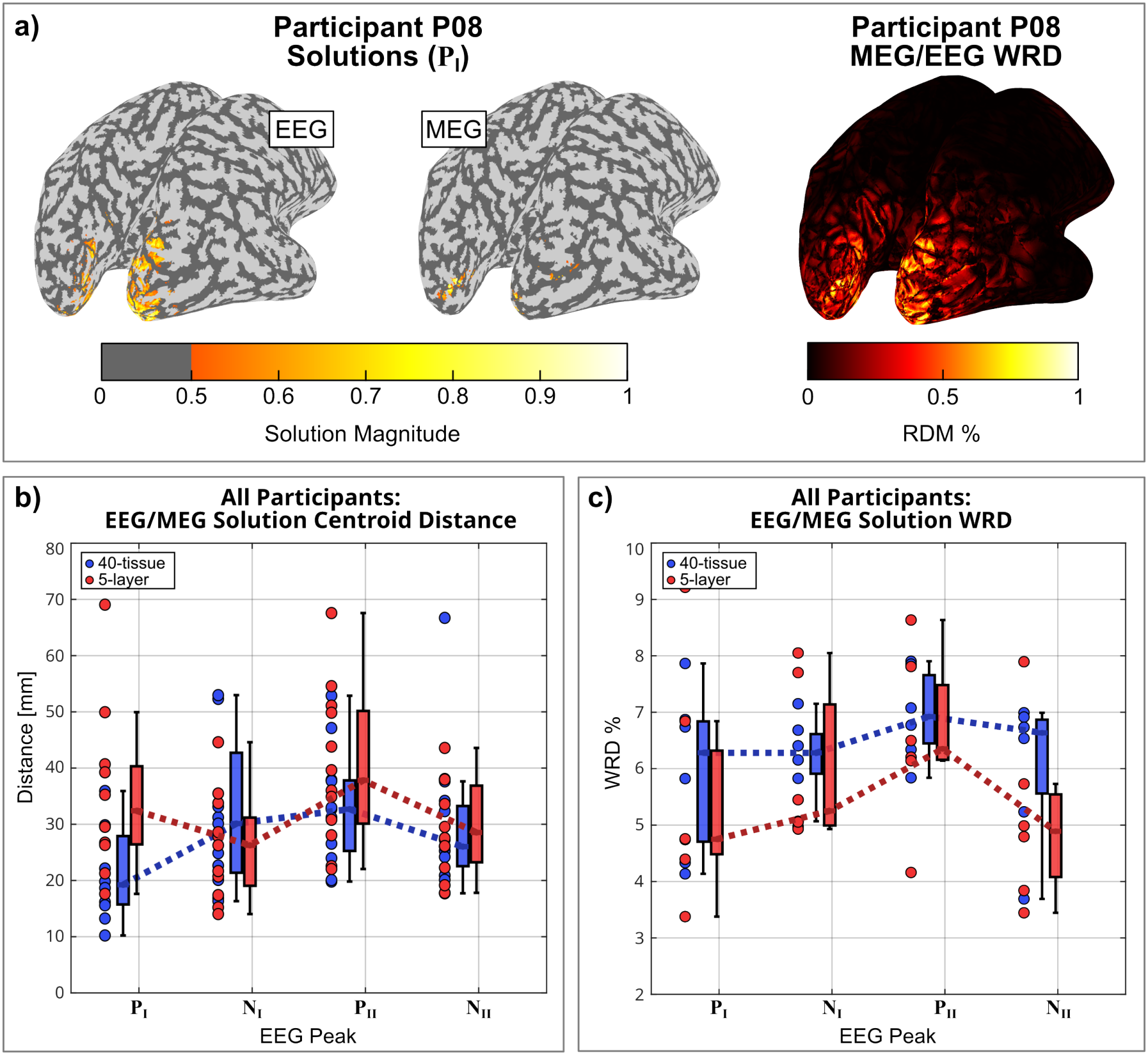
a) The source localization solutions for participant P08 (40-tissue model) at the P_I_ peak latency using EEG (left) and MEG (middle) data, and the corresponding WRD percentage between the two solutions (right). b) The distance between the per-hemisphere solution centroids of the EEG and MEG solutions across all participants and at all EEG peak latencies, with 40-tissue models in blue and 5-layer models in red. c) The subsequent WRD percentage computed across all participants and at all EEG peak latencies. Tables with data points corresponding to this figure may be found in Supplement 3.

**Figure 5:**
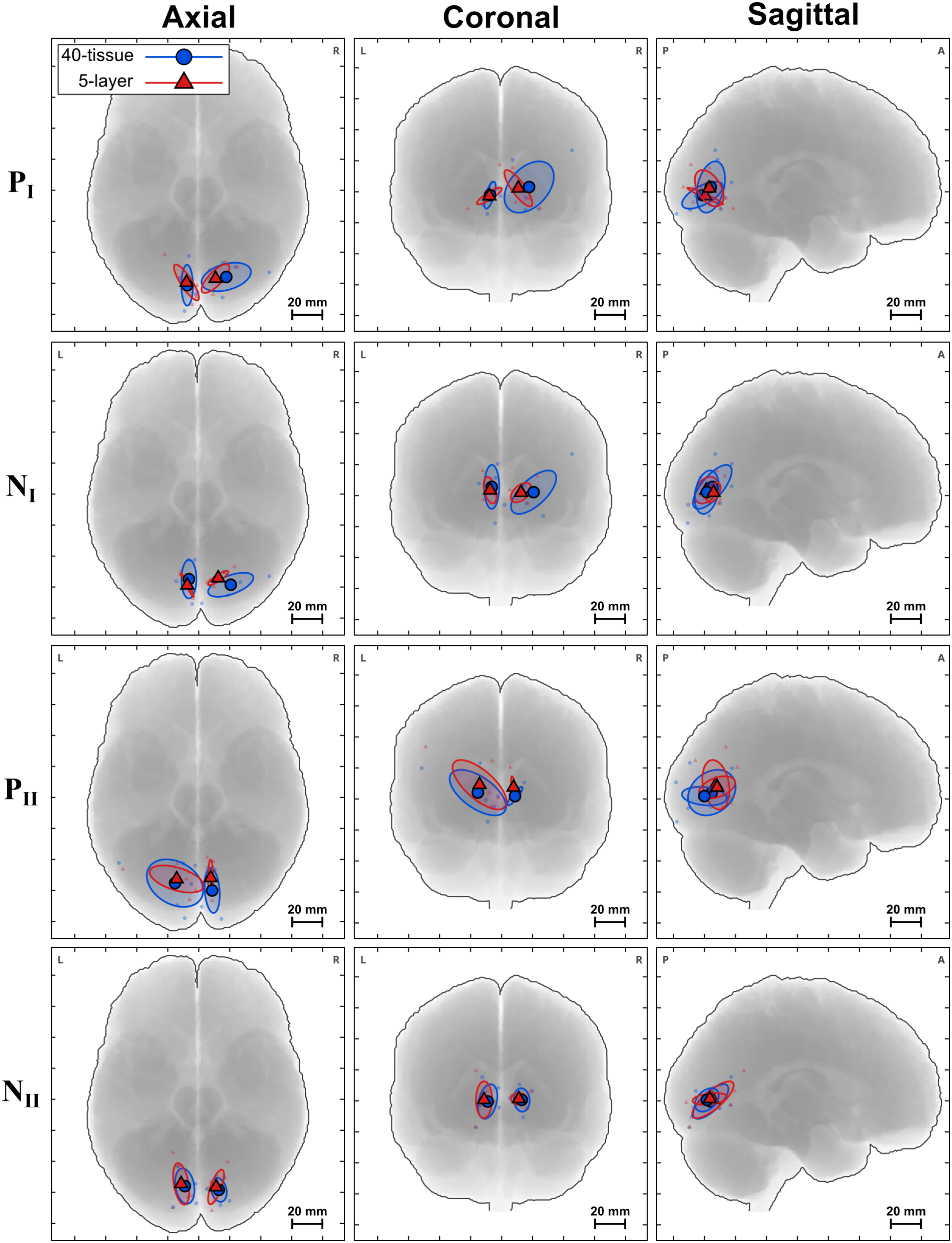
EEG source localization centroids of activation projected into the participant-averaged MNI space. Per-hemisphere solution centroids are shown for each of the four peaks, with the 40-tissue Sim4Life models in blue and the 5-layer SimNIBS model in red. Bold points with black outlines indicate the participant-averaged mean centroid, and the surrounding ellipses are regions *±* 1 standard deviation from the mean. Tables with data points corresponding to this figure may be found in Supplement 3.

### Synthetic leadfield analysis

The percentage of triangles processed according to our inclusion criterion was approximately 91% (90.8 *±* 0.7%). The median distance between each reference triangle and its closest headreco triangle after co-registration was 0.60 *±* 0.06mm (95th percentile, 1.53 *±* 0.20mm). The global scale factor between the 40-tissue and 5-layer models was 1.10 *±* 0.13 (range 0.86–1.26), i.e. the 5-layer leadfields were on average 10% larger. The raw median RDM between both models was 0.23 *±* 0.09, while the RDM obtained when comparing the 40-tissue model against itself, which sets the floor for neighboring locations of the same model, was 0.086 *±* 0.003. The XRDM, i.e. the raw RDM minus this baseline, was therefore 0.14 *±* 0.09. This cohort average is driven by a single participant (P10), whose median RDM was 0.41 (excess 0.32). In the remaining five participants, the median RDM was 0.196*±* 0.006 and the XRDM 0.11 *±* 0.01, about 2.3 times the baseline, which corresponds to a median topography correlation of approximately 0.98 between models. Regarding magnitude, the lnMAG* had a median of *−*0.004 *±* 0.003 and, for the central 50% of triangles, ranged from *−*0.09 to 0.09 (ratios of 0.91 to 1.09), about three times the width of the within-model baseline (0.18 vs. 0.06). In Figure 3 the comparison results for one participant (P03) are exhibited. Complete tables of leadfield comparison results can be found in Supplement 3.

## 4 Discussion

### Main findings

Our computational results show that EEG source localization with highly complex non-nested head models is feasible with our methodology. However, our source localization comparison results are inconclusive and cannot support the hypothesis that 40-tissue head models provide an advantage over simpler models. On the experimental side of our work, we have identified four EEG peaks in visual stimulation at one-half of the iAPF, with a fairly consistent waveform across all participants. This periodic waveform appears to correspond to the rhythmic component of the photic driving response. Hence, our analysis complements that of (Oppermann et al. 2025), where transient events were extracted and studied. Our leadfield comparison results show a nearly linear dependency between RDM and the angle between reference triangles and averaged centroids. This finding supports the hypothesis that an accurate alignment of the true surface normals to the dipole directions of the source space is very important in a source model employing fixed orientation constraints.

### Source localization

Although the small sample size prevents formal statistical hypothesis testing, we can draw some preliminary conclusions from our results. Upon visual inspection of the source localization solutions (see Figure 4a, and the corresponding figures in Supplement 2), there is no evidence that using the 40-tissue head model, rather than the 5-layer model, increases the focality of the EEG solutions compared to the MEG solutions. Rather, the MEG solutions are generally much more focal than the EEG solutions for both head models, which is perhaps expected given that there are more MEG sensors, and thus more independent measurements constraining the solution. Despite this visual difference, Figures 4b and 4c suggest that the EEG solutions from both head models agree with their corresponding MEG solutions to a similar degree. The average centroid distance between the EEG and MEG solutions lies between 20 and 40 mm across all peaks. Although this seems large, the entire visual cortex spans approximately 40 mm, so the EEG and MEG solutions generally fall within the same expected region. Further, the mean WRD is between 4 and 7% across all peaks for both models. The mean WRD is lower for the 5-layer model across all peaks, although this difference is probably negligible, as the cross-model comparisons below will show. That is, increasing the number of tissues does not significantly improve the agreement between sources localized from EEG and MEG data.

The direct comparisons between the 40-tissue and 5-layer EEG solutions in Table 2 support this. Across all participants, peaks, and hemispheres, the mean distance between the centroids of the two models is 12.3 mm, and 22 of the reported distances are below 10 mm. The distances are largest for P_I_ and N_I_ and smallest for P_II_ and N_II_, whereas the WRD is similar for all peaks, with a mean below 3%. The two models therefore produce very similar localization solutions overall (see also the figures in Supplement 2), and the larger centroid distances at some peaks reflect shifts of the most strongly activated region rather than differences across the whole map. When the centroids are aggregated across participants in MNI space (Figure 5), the two models agree for all peaks. The 5-layer centroids cluster more tightly across participants than the 40-tissue centroids. This may be due to the high inter-participant variability of the many non-nested tissues in the 40-tissue models compared with the simpler nested 5-layer geometry, although with six participants this difference in spread should be interpreted with caution. The agreement between the models is greatest for N_II_, which has the smallest centroid distances in Table 2 and the largest overlap in MNI space.

### Model leadfield difference

Given that our modeling pipeline is the same for both models, the fact that the global scale factor was near 1 on average indicates that the choice of surrogate conductivity (0.229S m^*−*1^) for the skin layer in the 5-layer model provides a good agreement with the 40-tissue models even without individualized depth weighing for conductivity assignment. Our leadfield comparisons show a median XRDM of 0.14 *±* 0.09 with respect to the baseline in coregistered regions of similar location and orientation. Although our baseline criterion is rather conservative, since the average edge length of the 40-tissue models (~ 1.5mm) is about 2.5 times larger than the co-registration distance between the 40-tissue and 5-layer models (~0.6mm), the raw RDM between the 40-tissue and 5-layer models is very small (0.23 *±* 0.09) in itself. This appears to indicate a fairly large agreement between both leadfield solutions. It must be mentioned, however, that due to our criterion of selecting only triangles within a small angle of the reference triangle, the majority of the regions being analyzed correspond to sulcal walls. This is because sulcal walls are the regions where the co-registration between both source spaces matches more closely, gyral crests appear to occupy a larger volume of space in the headreco segmentation as shown in Figure 1 a). Otherwise, due to the apparent uniform distribution of RDM throughout the cortex there is little evidence to support the hypothesis that there should be a larger RDM at the gyral crests or deep sulci, and thus it appears that the RDM difference is largely a global effect across the entire cortex. As Figure 3g) shows, there is an approximately linear relationship between RDM and the angle between reference sources and averaged centroids. This indicates that an accurate representation of the true source directions in the source space is of great importance. Since FEM cannot prescribe arbitrary dipole sources with arbitrary orientations, and is instead forced to approximate fields within the mesh elements, this result suggests that BEM-FMM with an accurate representation of surface normals could be preferred over FEM when a fixed orientation is enforced. In general, this result supports the use of loose, rather than fixed, orientation constraints in source localization, gauged in function of the smoothness of the source space normals. A systematic comparative analysis of the effect of loose orientation constraints in the 40-tissue model could be a good line of future research.

### Runtime

We point out that both the runtime and memory usage could possibly be reduced by the use of less neighbor integrals and a looser relative residual, as well as by an optimized implementation. We hypothesize that 512 neighbor integrals and a relative residual of 10^*−*3^ could be sufficient for the present application; but to draw formal conclusions a systematic study of forward modeling parameters against source localization results would be necessary. A formulation of BEM-FMM with linear, rather than zeroth-order, basis functions in the Galerkin approximation could achieve similar accuracy on coarser meshes, which would further reduce computational cost. Whether it would also reduce the number of neighbor integrals required for the near-field correction remains to be investigated.

### Validation and practical implications

Due to the small sample size in our cohort, the lack of ground truth data, and the scarcity of literature on visual stimulation in the frequency range we study here, we cannot establish a significant difference in the accuracy of source localization solutions between the 40-tissue and simpler 5-layer models. The fact that all signal peaks are localized within the visual cortex supports the overall correctness of the localization results. However, our results do not support the hypothesis that a finer model than the typical nested 5-layer model will provide better localization results for healthy participants. We note, however, that a detailed non-nested model may be very relevant for EEG localization in infants and people with pathologies.

This is because nested layered models cannot represent openings in the skull such as fontanels and sutures, which are particularly relevant in infants. Such features have been shown to distort EEG more strongly than MEG (Lew et al. 2013), as well as in patients with post-surgical skull defects (Lau et al. 2016). Likewise, lesions and tumors cannot be expressed using nested geometries. While a comparative analysis of solutions of a non-nested model with fontanels, sutures, skull defects, or tumors against the nested 3–5 layer models is out of the scope of this paper, we note that the methods presented here can be directly applied to head models incorporating these features. Such an analysis is an interesting topic of future research.

### Further future directions

A comparative analysis of the sensitivity of leadfields to the introduction of noise, or with respect to conductivity uncertainty, may reveal interesting information. Results showing that the 40-tissue models are less sensitive to noise or conductivity uncertainty may support their implementation into source localization pipelines. If such an analysis produces negative results, then it could steer source localization research away from including more tissue compartments in the forward modeling and instead focus more on accurately representing the direction of dipoles in the source space.

## Supporting information

Supplemental theory, figures, and tables

## Conflicts of Interest

The authors declare no conflicts of interest.

## Statement of AI use in research

Generative AI (Claude Sonnet 5.5 and Opus 5.5) was employed in code generation for the creation of figures and comparative analyses, as well as to improve the writing quality of the paper. All of the code was manually reviewed to ensure accuracy, and all authors have manually reviewed and edited the final version of the paper.

## Acknowledgments

G.N.P., D.A.D., G.M.N., and S.N.M. were supported by the NIBIB Grant 1R01EB035484, and the NIMH Grant 1R01MH130490. H.O. received funding from the Thuringian Ministry for Economic Affairs, Science and Digital Society within the “Learning Products” project (5575/10-3). B.M. was supported by the German Federal Ministry of Education and Research (BMBF) grant DryPole (01GQ2304B). J.H. received funding from the German Federal Ministry of Education and Research (BMBF) grant DryPole (01GQ2304A) and the Free State of Thuringia (2018 IZN 004), co-financed by the European Union under the European Regional Development Fund (ERDF).

## Notes

### Competing Interest Statement

The authors have declared no competing interest.

