## Supplemental theory, figures, and tables for "EEG Source Localization with High-Resolution 40-Tissue Head Models: A Comparative Study with MEG during Photic Driving": supp01_Theory.pdf

Friday 20th November, 2026

### 1 Derivation of Helmholtz Reciprocity for Current Electrodes and EEG

We derive in detail the EEG reciprocity relation for current electrodes and point dipoles. This is following and expanding upon the exposition of Plonsey 1963.

**Derivation of quasi-static boundary value problems in bioelectromaggnetism.** In bioelectromagnetism, the use of the quasi-static approximation to Maxwell's Equations is justified (Plonsey and Heppner 1967). In practice, this means that we may neglect all time derivatives.

Denote the volume conductor (human head) by $\Omega$ and its surface by $\partial\Omega$. We will consider two types of problems, one with source currents only in the interior $\mathring{\Omega}$ (case 1) of the head and another with source currents only on the surface $\partial\Omega$ (case 2) of the head.

**Case 1. Sources only in the interior of the head (EEG forward).** Let $\mathbf{J}_1^i$ be a primary, or impressed, current density which is zero. The total current $\mathbf{J}_1$ is split into two components

$$\mathbf{J}_1 = \mathbf{J}_1^i + \mathbf{J}_1^s, \tag{1}$$

where $\mathbf{J}_1^s$ is the secondary current, or volume current. In the case of EEG, the primary current can be approximated by a mathematical point-dipole in free space at a location $\mathbf{p}$ (de Munck et al. 1988):

$$\mathbf{J}_1^i(\mathbf{r}) = \mathbf{q} \cdot \delta(\mathbf{r} - \mathbf{p}), \tag{2}$$

where $\mathbf{q}$ is the dipole moment. In particular, the primary field is everywhere continuous except at the singular point $\mathbf{p}$. By Ohm's Law $\mathbf{J}_1^s = \sigma \mathbf{E}_1^s$, where $\sigma$ can be regarded as a 3-tensor $\mathbb{R}^3 \to \mathbb{R}^3 \times \mathbb{R}^3$, and $\mathbf{E}_1^s$ is the volume electric field, i.e. the electric field elicited by the deposited charge density in the presence of the primary current $\mathbf{J}_1^i$. Namely, $\mathbf{E}_1^s$ is obtained from a charge density $\rho$ via Coulomb's Law, and as such this field is conservative and we can write $\mathbf{E}_1^s = -\nabla\phi_1$ for some scalar electric potential $\phi_1$.

Applying the quasi-static approximation to the continuity equation yields:

$$\nabla \cdot \mathbf{J}_1 = -\frac{\partial \rho}{\partial t} = 0. \tag{3}$$

Combining Equations 1 and 3, we arrive at the fundamental equation:

$$\nabla \cdot \sigma \nabla \phi_1 = \nabla \cdot \mathbf{J}_1^i; \tag{4}$$

which must be satisfied everywhere in the interior of $\Omega$.

A boundary condition can be also obtained from the continuity equation. First, note that we may approximate the conductivity of air as 0, so the skin surface $S$ is a discontinuity surface for $\sigma$. Consider a Gaussian pillbox volume $V$ centered at a point $\mathbf{r} \in S$, then the integral form of the continuity equation implies:

$$0 = \int_V \nabla \cdot \mathbf{J}_1 dv = \int_{\partial V} \mathbf{J}_1 \cdot \mathbf{n} ds. \tag{5}$$

Letting the thickness of the pillbox approach 0, we find

$$0 = (\mathbf{J}_1)_+ \cdot \mathbf{n} - (\mathbf{J}_1)_- \cdot \mathbf{n}, \tag{6}$$

where $(\mathbf{J}_1)_\pm$ denotes the limiting values of $\mathbf{J}_1$ just outside and just inside of the skin surface $S$. In other words, the normal component of the total current is continuous across the surface $\partial\Omega$. We assumed that there are no impressed currents on the surface $\partial\Omega$, and there are no volume current outside of $\Omega$, therefore $(\mathbf{J}_1)_+ = 0$ and we arrive at the boundary condition

$$\sigma \nabla \phi_1 = 0. \tag{7}$$

Putting everything together, we arrive at the following boundary value problem (which in particular governs EEG forward solutions):

$$\begin{cases} \nabla \cdot \sigma \nabla \phi_1 = \nabla \cdot \mathbf{J}_1^i & \text{in } \Omega \\ \sigma \nabla \phi_1 \cdot \mathbf{n} = 0 & \text{on } \partial\Omega \end{cases}, \tag{8}$$

where $\phi_1$ is the scalar potential of the conservative field induced by the charge density in $\Omega$ in presence of the impressed current $\mathbf{J}_1^i$.

**Case 2. Impressed sources only on the head surface (TES forward).** We can replicate the same arguments as above, now with an impressed current $\mathbf{J}_2^i$ which is zero everywhere in the interior of $\Omega$ and non-zero only at some parts of the skin surface $\partial\Omega$. For example, in the case of TES we may take the following impressed current density

$$\mathbf{J}_2^i(\mathbf{r}) = \delta(\mathbf{r} - \mathbf{r}_1) - \delta(\mathbf{r} - \mathbf{r}_2), \tag{9}$$

where $\mathbf{r}_1$ and $\mathbf{r}_2$ are the centers of two voltage electrodes on the skin surface.

The only difference is that now, $\mathbf{J}_2^i = 0$ in $\Omega$ so the differential equation in the interior is

$$\nabla \cdot \sigma \nabla \phi_2 = 0 \quad \text{in } \Omega. \tag{10}$$

And on the surface we have that the total current just outside is precisely given by the impressed current $\mathbf{J}_2^i$ (the current impressed by the electrodes on the skin surface). Hence, we obtain the boundary condition

$$\sigma \nabla \phi_2 \cdot \mathbf{n} = \mathbf{J}_2^i \quad \text{on } \partial\Omega. \tag{11}$$

Which put together reads as the following boundary value problem:

$$\begin{cases} \nabla \cdot \sigma \nabla \phi_2 = 0 & \text{in } \Omega \\ -\sigma \nabla \phi_2 \cdot \mathbf{n} = \mathbf{J}_2^i & \text{on } \partial\Omega \end{cases}, \tag{12}$$

where $\phi_2$ is the scalar potential of the conservative field induced by the charge density in $\Omega$ in presence of the impressed current $\mathbf{J}_2^i$. Here, we are assuming that the normals of $\partial\Omega$ are directed inwards (hence the negative sign appearing).

**Green's Theorem.** We have the following identities for the "cross-terms" of the two potentials $\phi_1$ and $\phi_2$ described above:

$$\nabla \cdot (\phi_1 \sigma \nabla \phi_2) = \sigma \nabla \phi_1 \cdot \nabla \phi_2 + \phi_1 \nabla \cdot (\sigma \nabla \phi_2) \tag{13}$$

$$\nabla \cdot (\phi_2 \sigma \nabla \phi_1) = \sigma \nabla \phi_2 \cdot \nabla \phi_1 + \phi_2 \nabla \cdot (\sigma \nabla \phi_1) \tag{14}$$

Applying Gauss's Divergence Theorem, and subtracting we obtain

$$\begin{aligned} \int_{\partial\Omega} \phi_1 \sigma \nabla \phi_2 \cdot \mathbf{n} ds - \int_{\partial\Omega} \phi_2 \sigma \nabla \phi_1 \cdot \mathbf{n} ds = \\ \int_{\Omega} \phi_1 \nabla \cdot (\sigma \nabla \phi_2) dv - \int_{\Omega} \phi_2 \nabla \cdot (\sigma \nabla \phi_1) dv. \end{aligned} \tag{15}$$

Now, applying the conditions in Equations 8 and 12 to $\phi_1$ and $\phi_2$ respectively, we find

$$\int_{\partial\Omega} \phi_1 \mathbf{J}_2^i \cdot \mathbf{n} ds = \int_{\Omega} \phi_2 \mathbf{J}_1^i dv. \tag{16}$$

This is the continuous version of Green's Theorem.

**EEG-TES reciprocity.** We now let

$$\mathbf{J}_1^i(\mathbf{r}) = \frac{1}{\varepsilon}\left[\delta(\mathbf{r} - (\mathbf{p} + \varepsilon\mathbf{d})) - \delta(\mathbf{r} - \mathbf{p})\right]; \tag{17}$$

here $\varepsilon$ is a dimensionless scaling constant. This is the impressed current density of a unit-strength finite length current dipole with displacement direction $\mathbf{d}$ (a unit vector) and length $\varepsilon > 0$.

On the other hand, we let

$$\mathbf{J}_2^i(\mathbf{r}) = \delta(\mathbf{r} - \mathbf{r}_k) + \delta(\mathbf{r} - \mathbf{r}_0), \tag{18}$$

where $\mathbf{r}_k$ is the center of the $k$-th electrode of an EEG sensor sytem, and $\mathbf{r}_0$ is the reference electrode. Substituting into Equation 16, we find

$$\phi_1(\mathbf{r}_k) - \phi_1(\mathbf{r}_0) = \frac{\phi_2(\mathbf{p} + \varepsilon\mathbf{d}) - \phi_2(\mathbf{p})}{\varepsilon}, \tag{19}$$

which holds true for all $\varepsilon > 0$. Note that this is an identity only involving potential differences, so it is consistent in terms of units. Now, using the definition of the derivative, we have

$$-\mathbf{E}_2^s(\mathbf{p}) \cdot \mathbf{d} = \nabla\phi_2 \cdot \mathbf{d} = \lim_{\varepsilon \to 0} = \frac{\phi_2(\mathbf{p}) - \phi_2(\mathbf{p} + \varepsilon\mathbf{d})}{\varepsilon} = \phi_1(\mathbf{r}_k) - \phi_1(\mathbf{r}_0). \tag{20}$$

The leftmost term is in units $[\mathrm{V\,m^{-1}} \cdot \mathrm{m}]$=$[\mathrm{V}]$, so this equation is also consistent in terms of units. We arrive at the fundamental Helmholtz's reciprocity relating EEG forward solutions with TES forward solutions:

$$-\mathbf{E}_2^s(\mathbf{p}) \cdot \mathbf{d} = \phi_1(\mathbf{r}_k) - \phi_1(\mathbf{r}_0), \tag{21}$$

where $\mathbf{E}_2^s$ is the volume field elicited by TES at cathode position $\mathbf{r}_k$ and anode position $\mathbf{r}_0$, and $\phi_1$ is the potential of the volume field elicited by EEG with a point dipole of unit direction $\mathbf{d}$ located at position $\mathbf{p}$.

### 2 EEG Reciprocity from Circuit Reciprocity

The quasistatic reciprocity (duality) principle or theorem (Helmholtz 1853; Rush and Driscoll 1969) makes it possible to interrelate the fields within the head generated by surface electrodes with the fields of the forward EEG problem with embedded dipolar point current sources (Vallaghé et al. 2008; Wagner et al. 2016). Also, the corresponding powerful analytical EEG solutions — cf. (Zhang 1995; Mosher et al. 1999) — could be used to test the validity of the fields injected by surface electrodes. In the relevant EEG studies (Wagner et al. 2016; Vanrumste et al. 2001; Laarne et al. 2000; Finke et al. 2013; Fletcher et al. 1995; Shirvany et al. 2013) the reciprocal solution was primarily used to more easily obtain multiple forward solutions with the elementary dipoles. Then, these forward solutions were employed in a standard framework of the EEG inverse analysis. We apply a somewhat different approach. It effectively bypasses the forward EEG solutions and utilizes the fields generated by surface electrodes for the solution of the inverse EEG problem over the entire cortical surface.

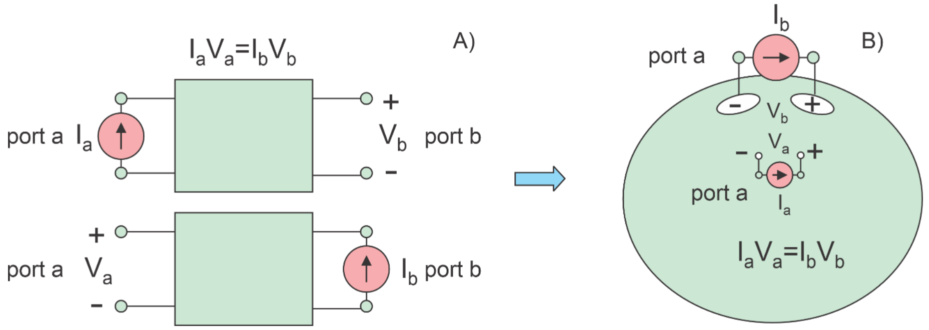


Figure 1: A. Circuital reciprocity theorem. B. Its application to a distributed circuit – a volume conductor.

**One electrode pair and one cortical dipole.** Among other methods, a straightforward way to derive the necessary relation is to directly apply the circuital reciprocity theorem (Lorrain et al. 1987; Guillemin 1953) for a linear passive circuit. The circuit is connected to a current source and a voltmeter as in Fig. 1A. In this case, interchanging the ideal current source and the voltmeter does not alter the ratio V/I so that $V_aI_a = V_bI_b$ (Lorrain et al. 1987; Guillemin 1953).

Nothing changes if the linear circuit takes the form of a volume conductor with many distributed resistances (Fig. 1B). We set Port a (a dipole current source) somewhere inside the conductor while Port b (a pair of surface current electrodes) will be located at its surface. We introduce the following notations: $I_a = I_d$ where $I_d$ is the dipole current strength, $V_a = -d \cdot E$ where $d$ is the dipole vector length and $E$ is the electric field generated by the surface current source at the dipole position, $I_b = I$ is the strength of the surface current source, and $V_b = v$ is the voltage between the surface electrodes that is generated by the dipole source. In terms of these notations, the equality $V_aI_a = V_bI_b$ becomes

$$-I_d(d \cdot E) = Iv. \tag{22}$$

**One electrode pair and multiple cortical dipoles.** We keep the surface current source in Fig. 1B *the same* but try out different internal dipole sources at different locations $r_n, n = 1, \ldots, N$. In other words, we turn these cortical dipoles on, one by one, and obtain, from Eq. 22

$$-I_{d_1}(d_1 \cdot E(r_1)) = Iv_1, -I_{d_2}(d_2 \cdot E(r_2)) = Iv_2, \ldots, -I_{d_N}(d_N \cdot E(r_N)) = Iv_N \tag{23}$$

Adding up all Eqs. 23 yields

$$-\sum_{n=1}^{N} I_{d_n}(d_n \cdot E(r_n)) = IV, \quad V = \sum_{n=1}^{N} v_n, \tag{24}$$

where $V$ is the total surface electrode voltage generated by the dipole layer when all dipoles are turned on. The continuous (integral) version of Eq. 24 has the form

$$-\int_S E(r) \cdot q(r) ds = IV. \tag{25}$$

Here, $q(r)$ $(A \cdot m/m^2)$ is the current dipole moment per unit cross sectional area of a continuous dipole layer with surface $S$, $E(r)$ (V/m) is the continuous field of the surface current source across the dipole layer, $I$ (A) is the strength of the surface current source and $V$ (V) is the voltage between the two surface electrodes generated by the dipole layer.

The reciprocity principle is only applicable to a single surface electrode pair at a time. All other electrode pairs, i.e., all other surface current sources must be turned off.
