## Supplemental theory, figures, and tables for "EEG Source Localization with High-Resolution 40-Tissue Head Models: A Comparative Study with MEG during Photic Driving": supp02_Figures.pdf

### EEG Source Localization with High-Resolution 40-tissue head models: A Comparative Study with Visual Evoked Potentials: Supplement 2

#### Supplemental Figures

Guillermo Núñez Ponasso[1*], Derek A. Drumm[1], Hannes Oppermann[2], Gregory M. Noetscher[1], Burkhard Maess[3], Thomas R. Knösche[3], Sergey N. Makaroff[1,4†], and Jens Haueisen[2†]

[†]Last two authors share senior authorship.
[1]Dept. of Electrical & Computer Engineering, Worcester Polytechnic Institute, Worcester, MA, USA
[2]Technische Universität Ilmenau, Ilmenau, Thuringia, Germany
[3]Max Planck Institute for Human Cognitive and Brain Sciences, Leipzig, Saxony, Germany
[4]Athinoula A. Martinos Center for Biomedical Imaging, Department of Radiology, Massachusetts General Hospital, Boston, MA, USA

November 17, 2026

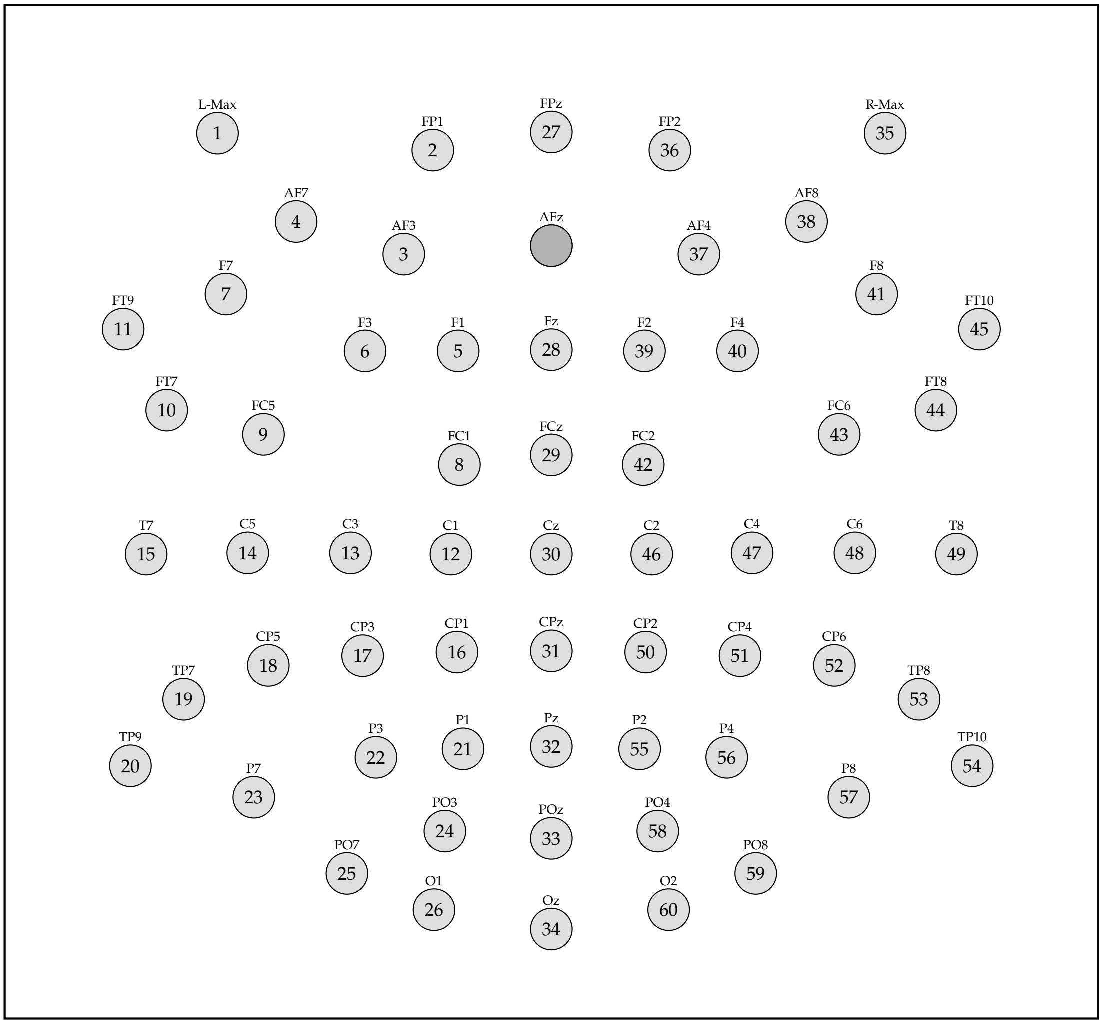


Figure S2.1: Layout of the 61-channel (waveguard™ CA-169, ANT b.v., The Netherlands) electrode cap.

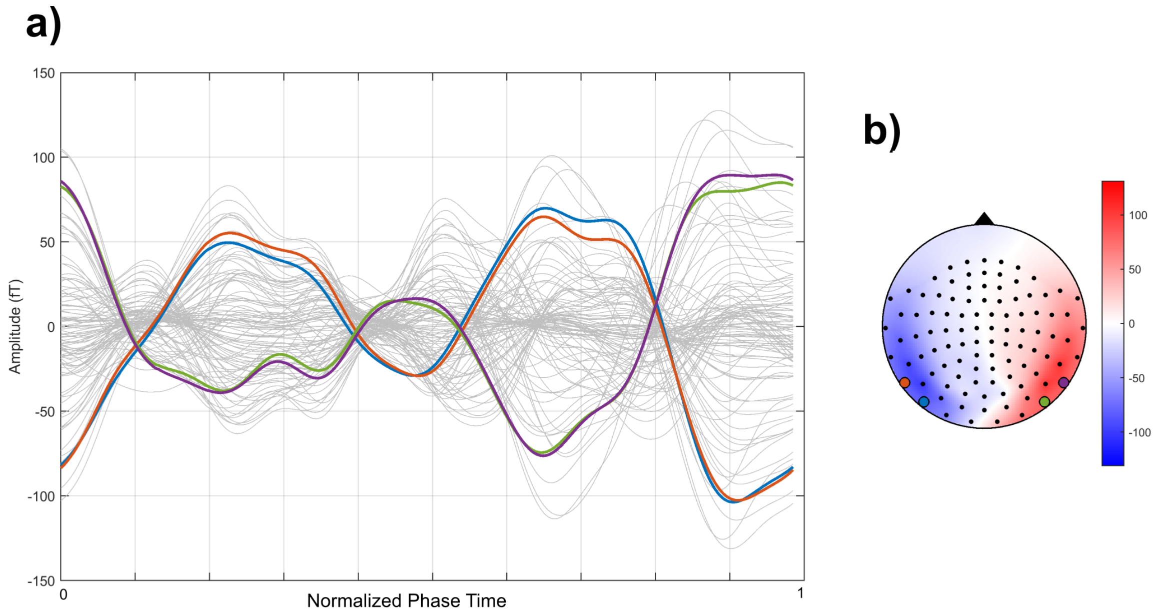


Figure S2.2: a) Participant-averaged MEG magnetometer signals plotted in phase time (with respect to EEG P1 peaks). Selected magnetometer sensor signals are highlighted. b) Corresponding topography with selected sensors highlighted.

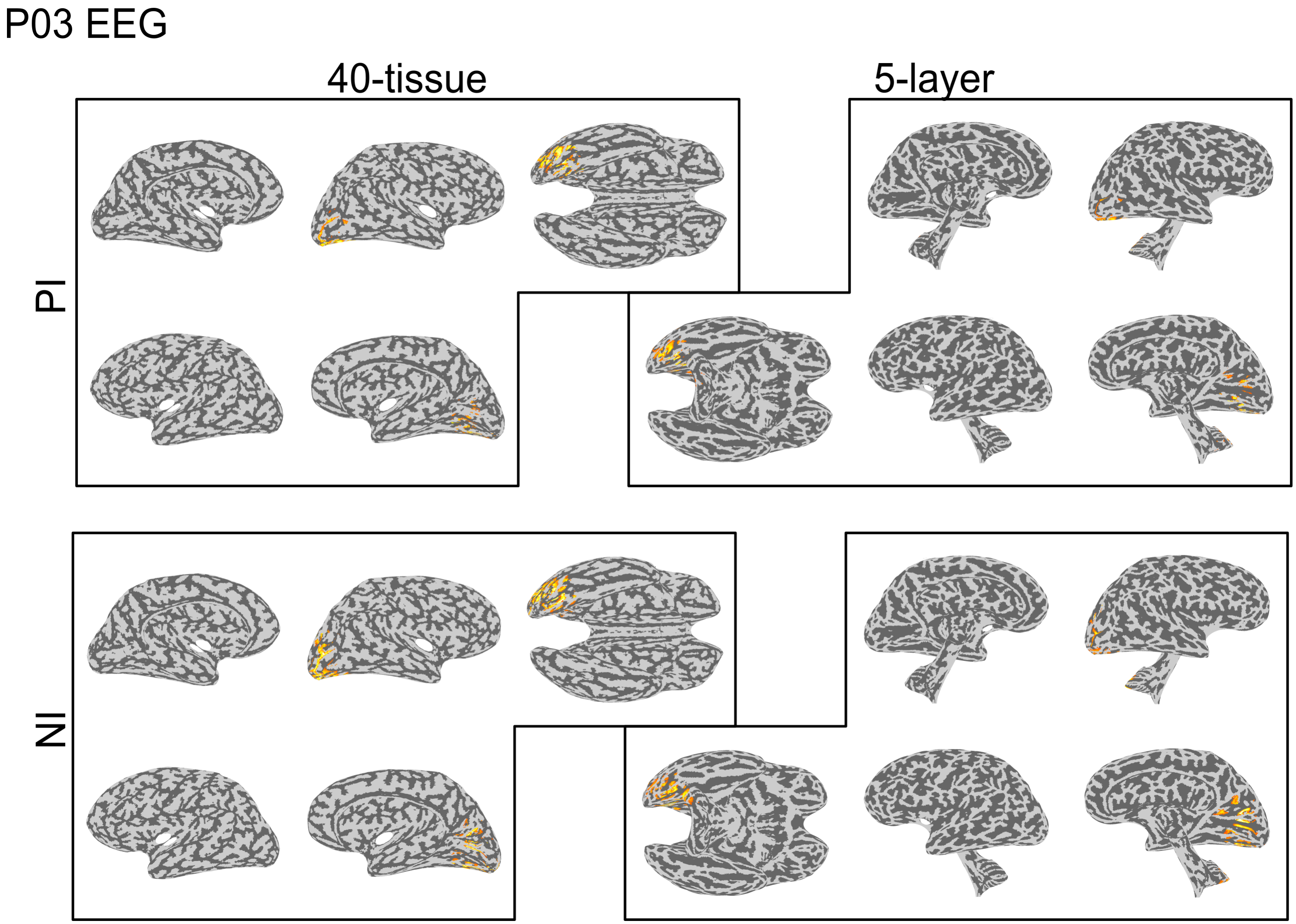


Figure S2.3: The EEG source localization solutions for subject P03 at each of the signal peaks ($P_I$ and $N_I$).

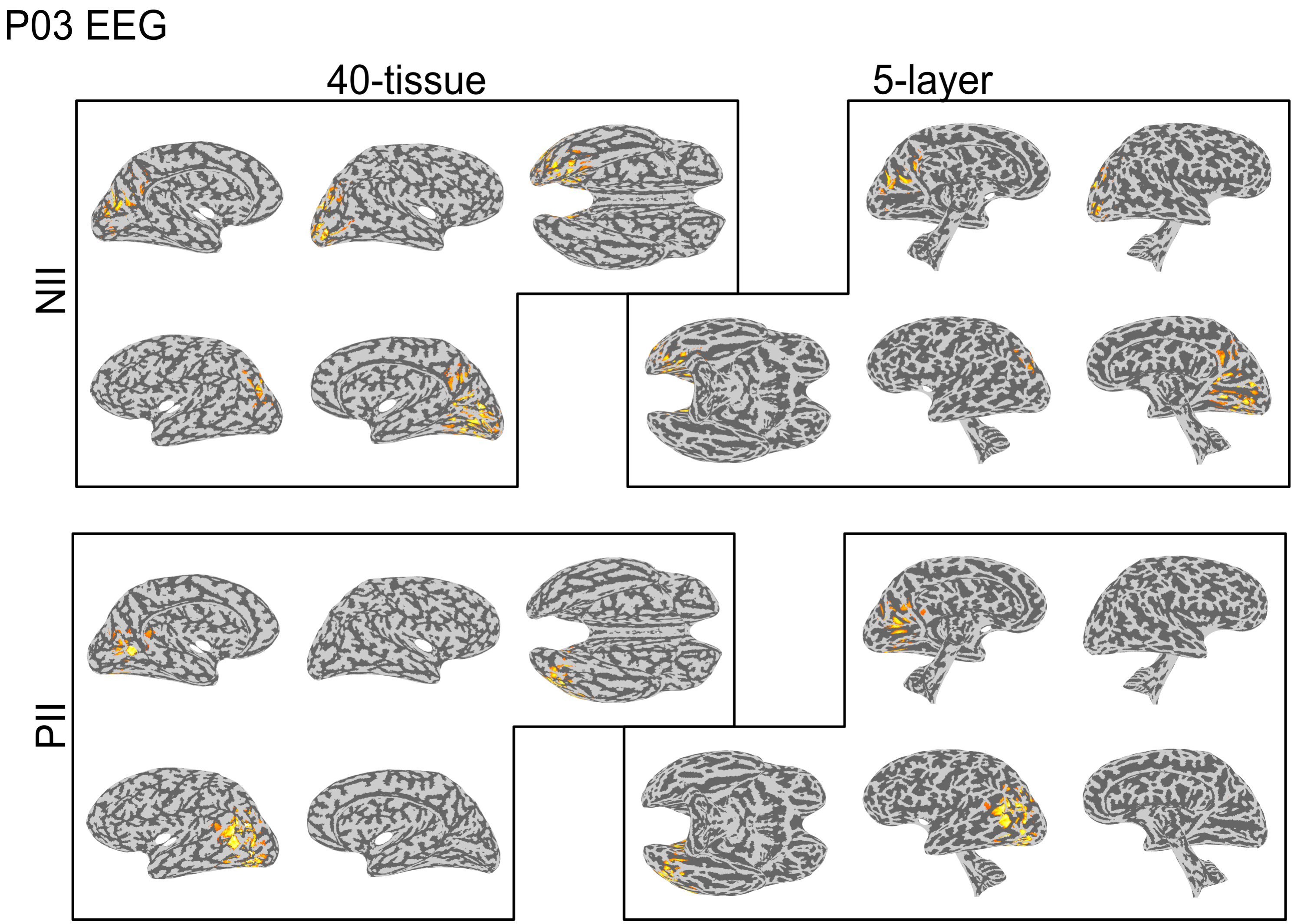


Figure S2.4: The EEG source localization solutions for subject P03 at each of the signal peaks ($N_{II}$ and $P_{II}$).

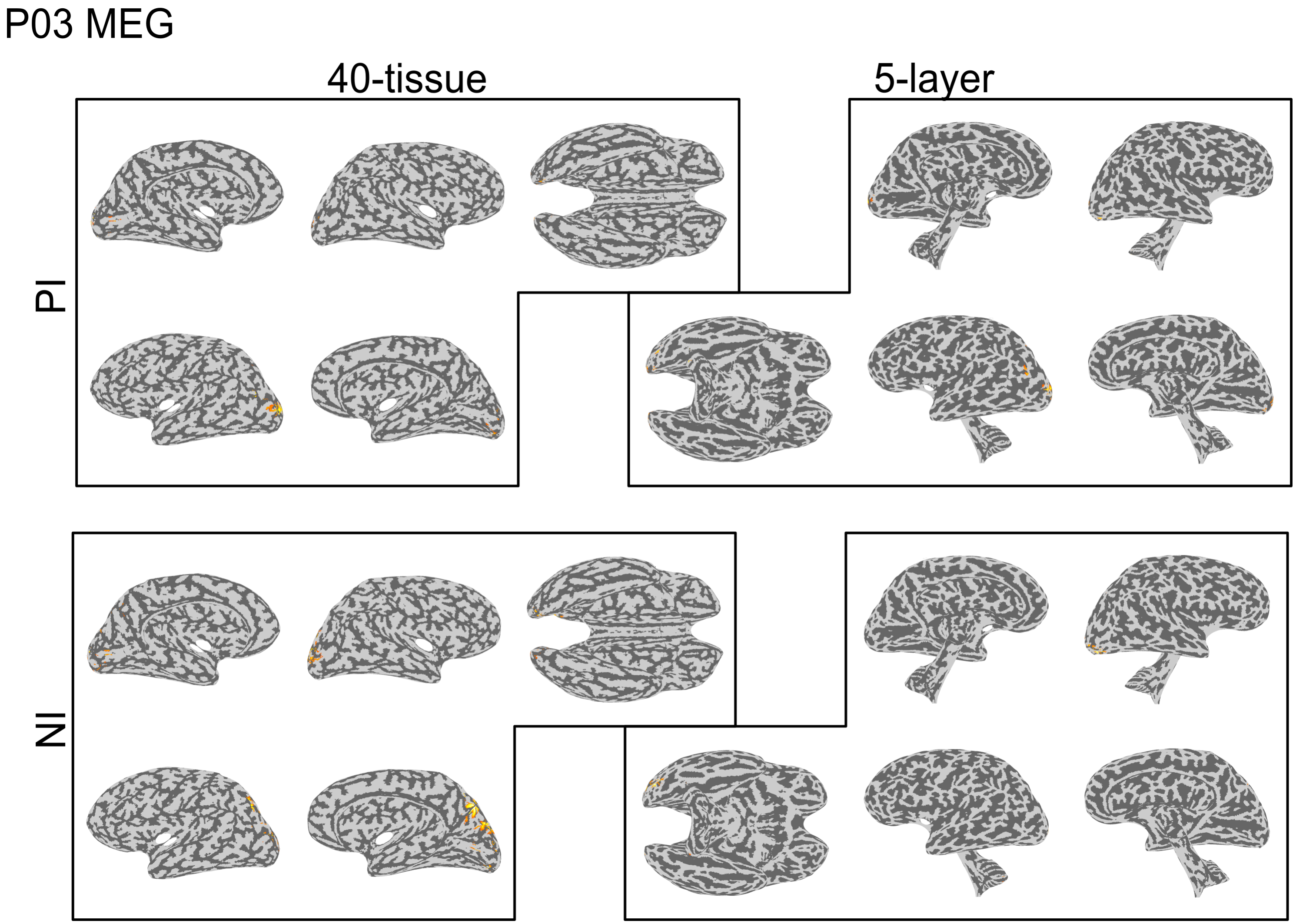


Figure S2.5: The MEG source localization solutions for subject P03 at each of the signal peaks ($P_I$ and $N_I$).

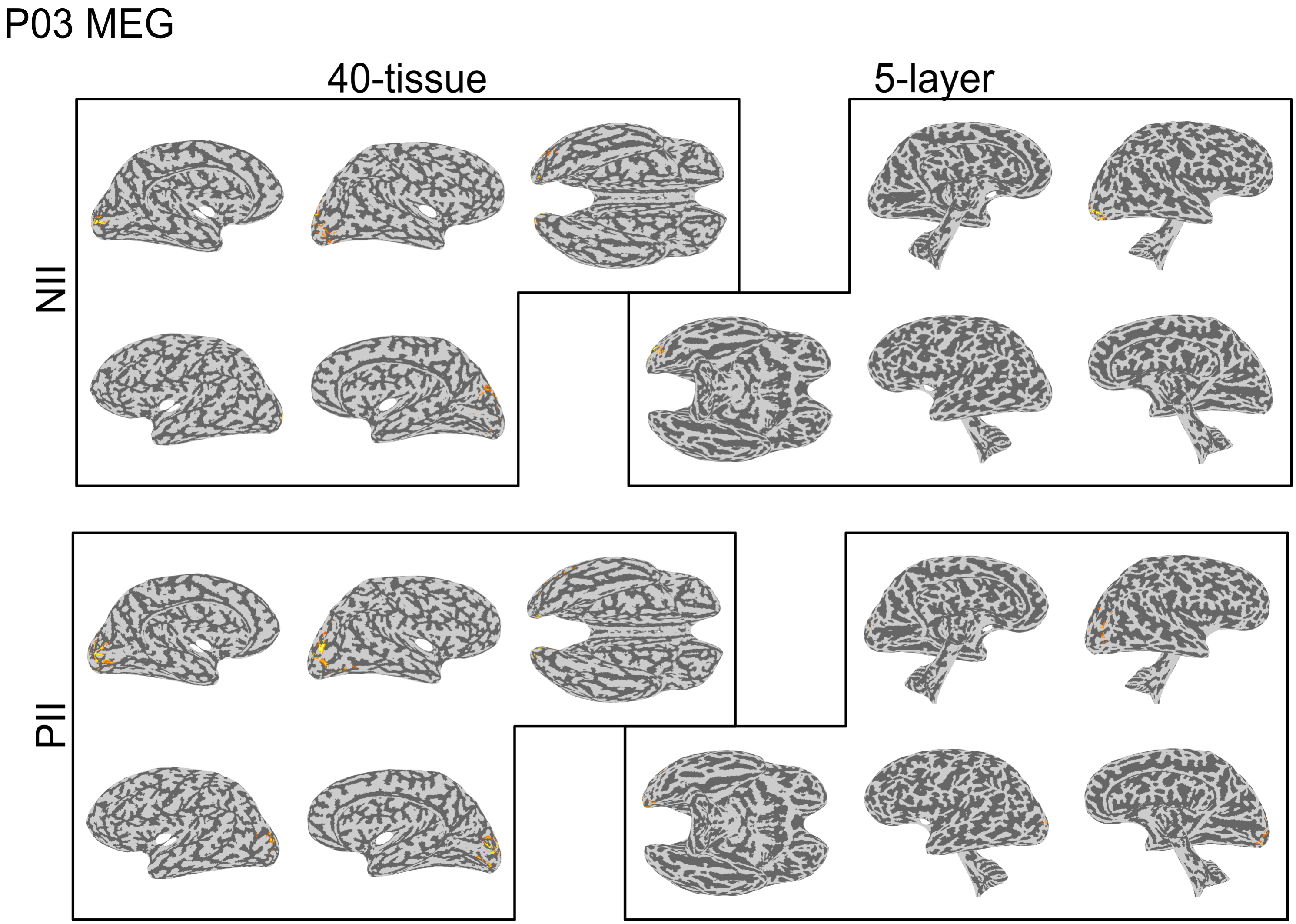


Figure S2.6: The MEG source localization solutions for subject P03 at each of the signal peaks ($N_{II}$ and $P_{II}$).

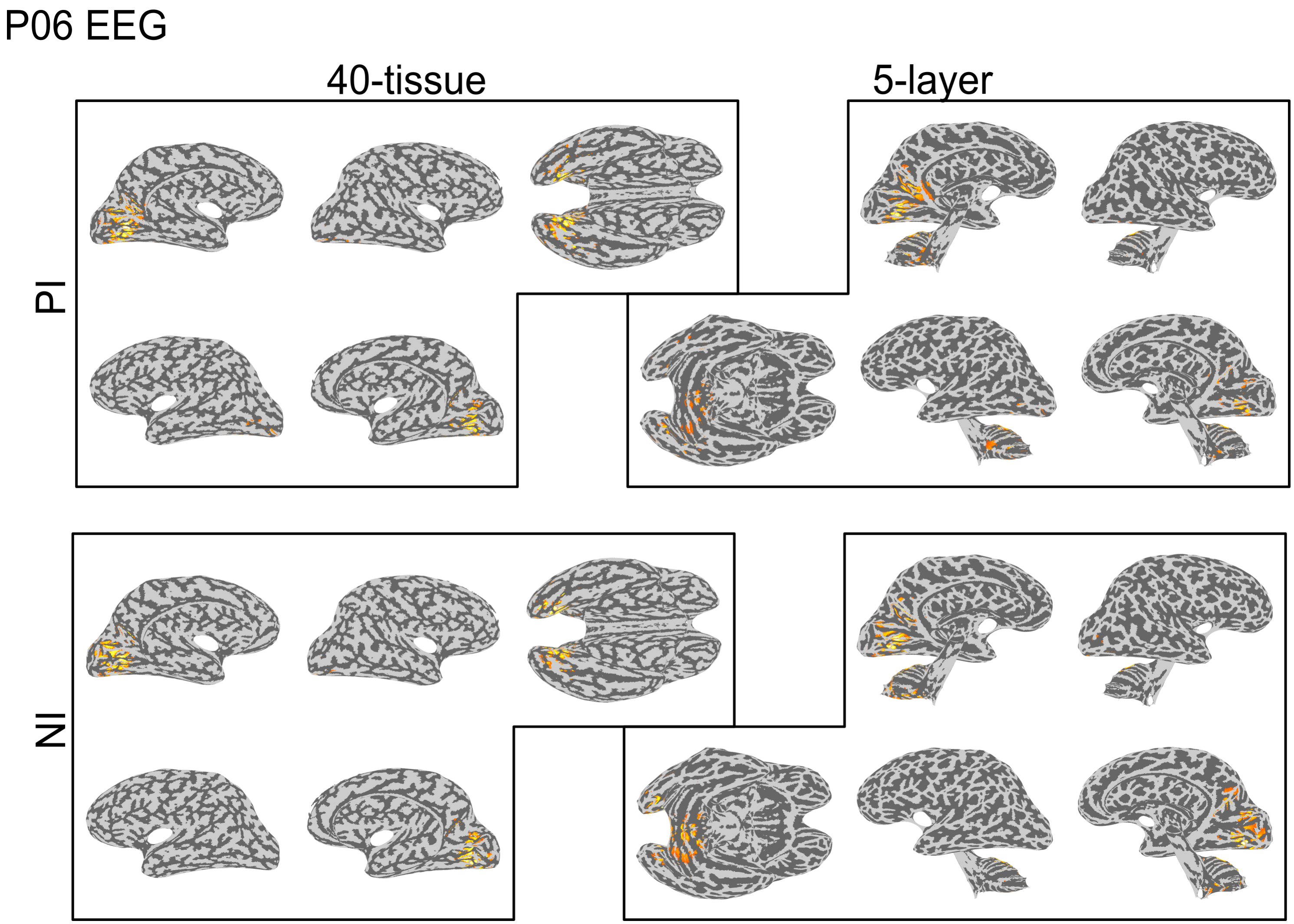


Figure S2.7: The EEG source localization solutions for subject P06 at each of the signal peaks ($P_I$ and $N_I$).

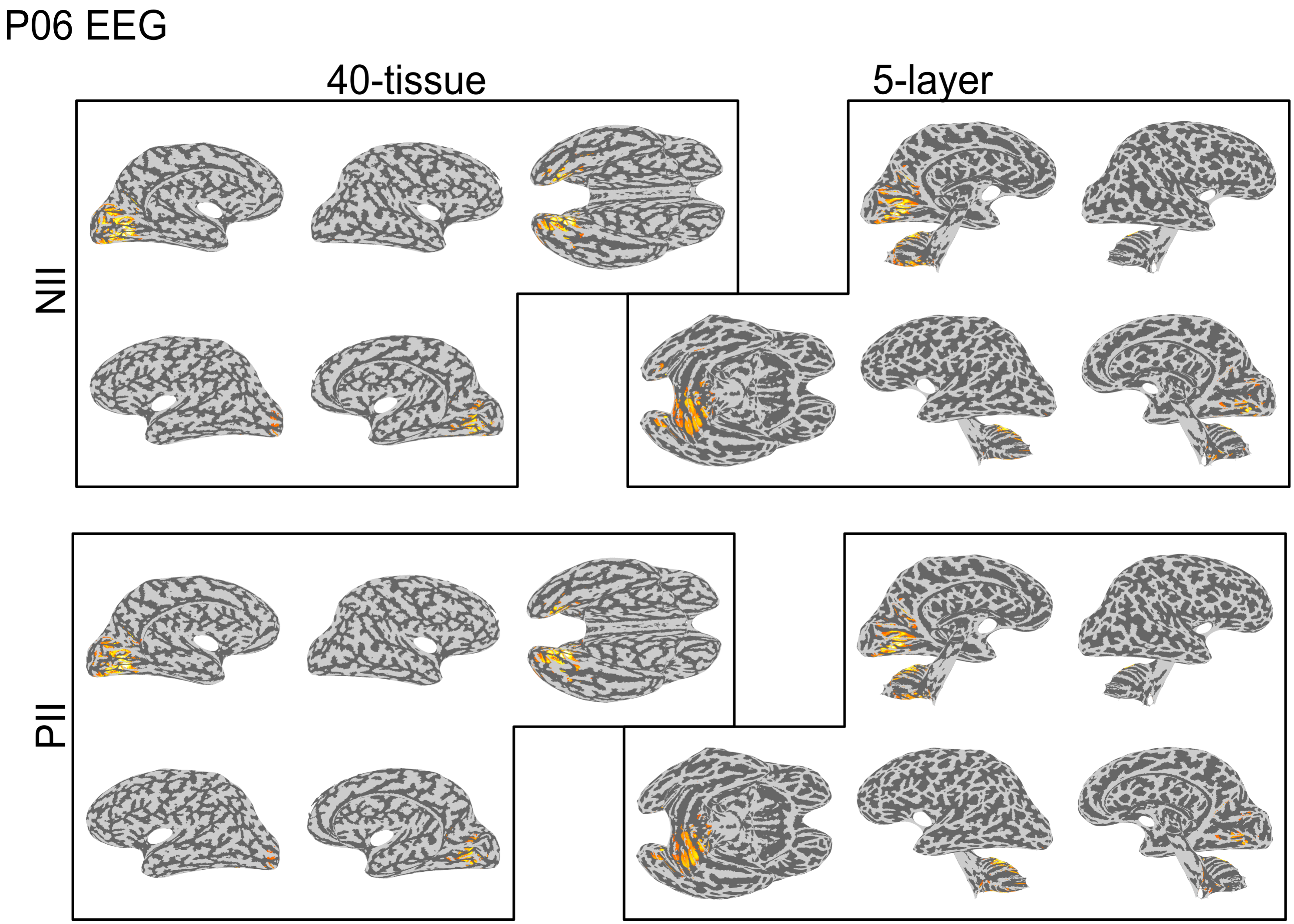


Figure S2.8: The EEG source localization solutions for subject P06 at each of the signal peaks ($N_{II}$ and $P_{II}$).

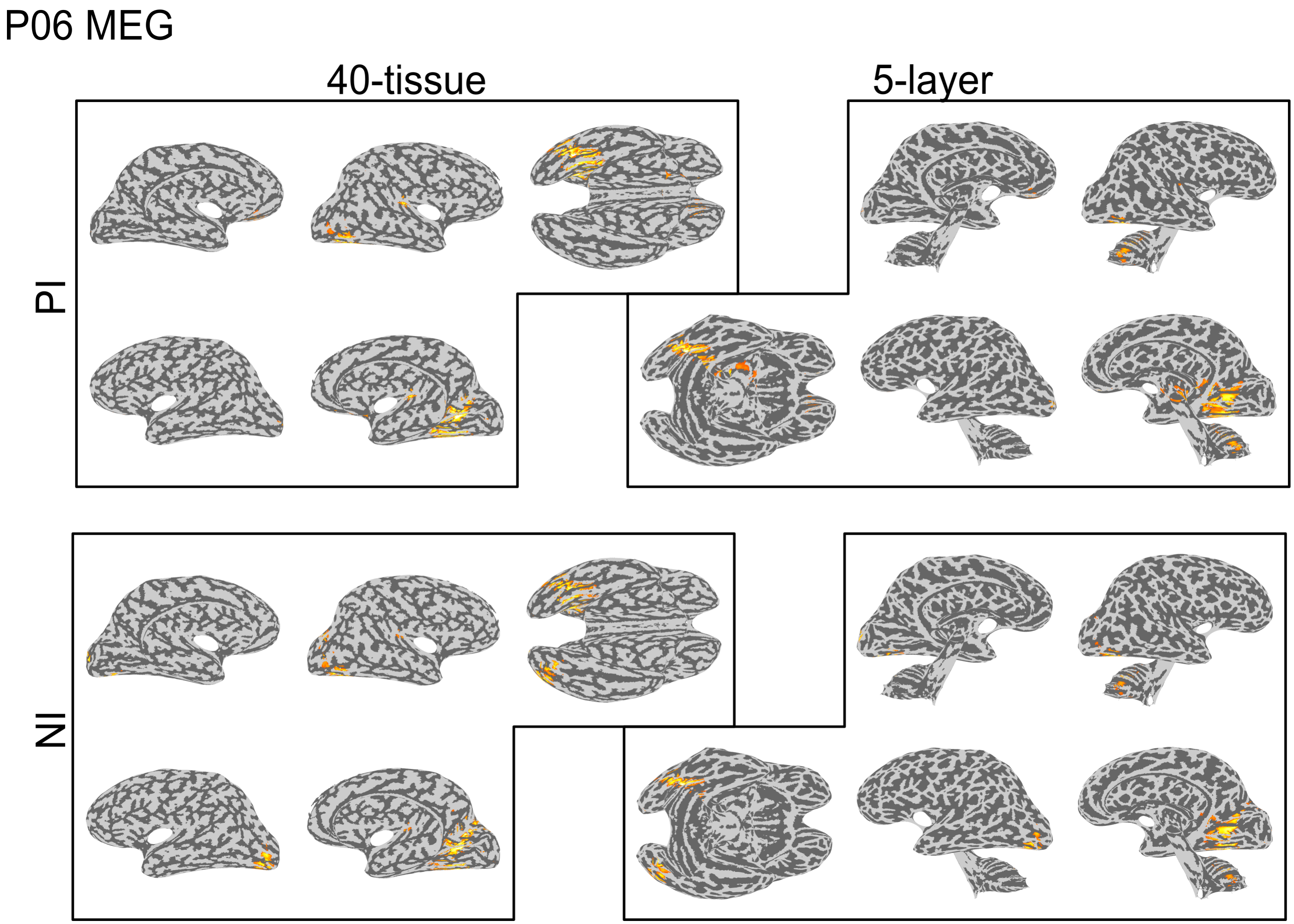


Figure S2.9: The MEG source localization solutions for subject P06 at each of the signal peaks ($P_I$ and $N_I$).

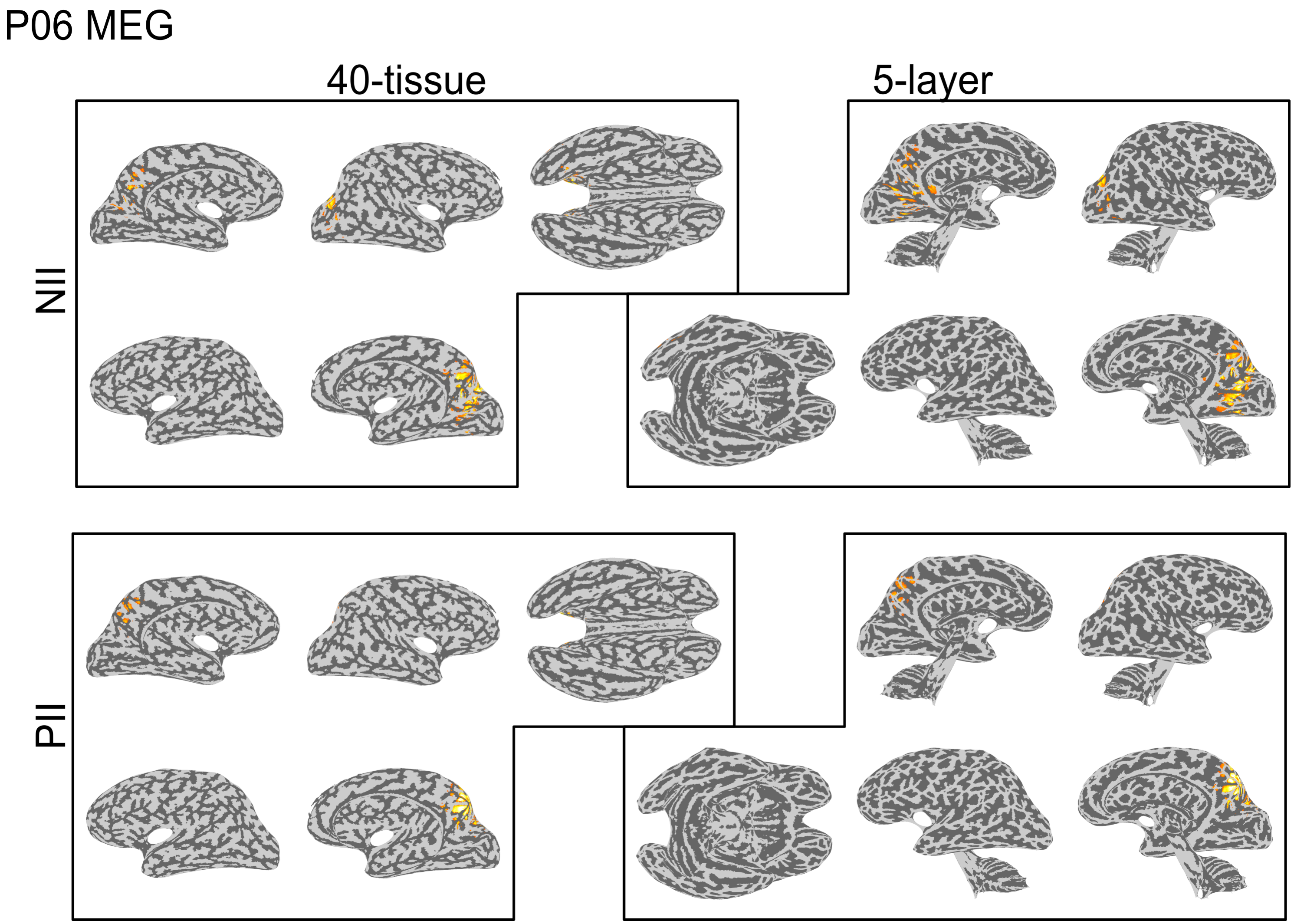


Figure S2.10: The MEG source localization solutions for subject P06 at each of the signal peaks ($N_{II}$ and $P_{II}$).

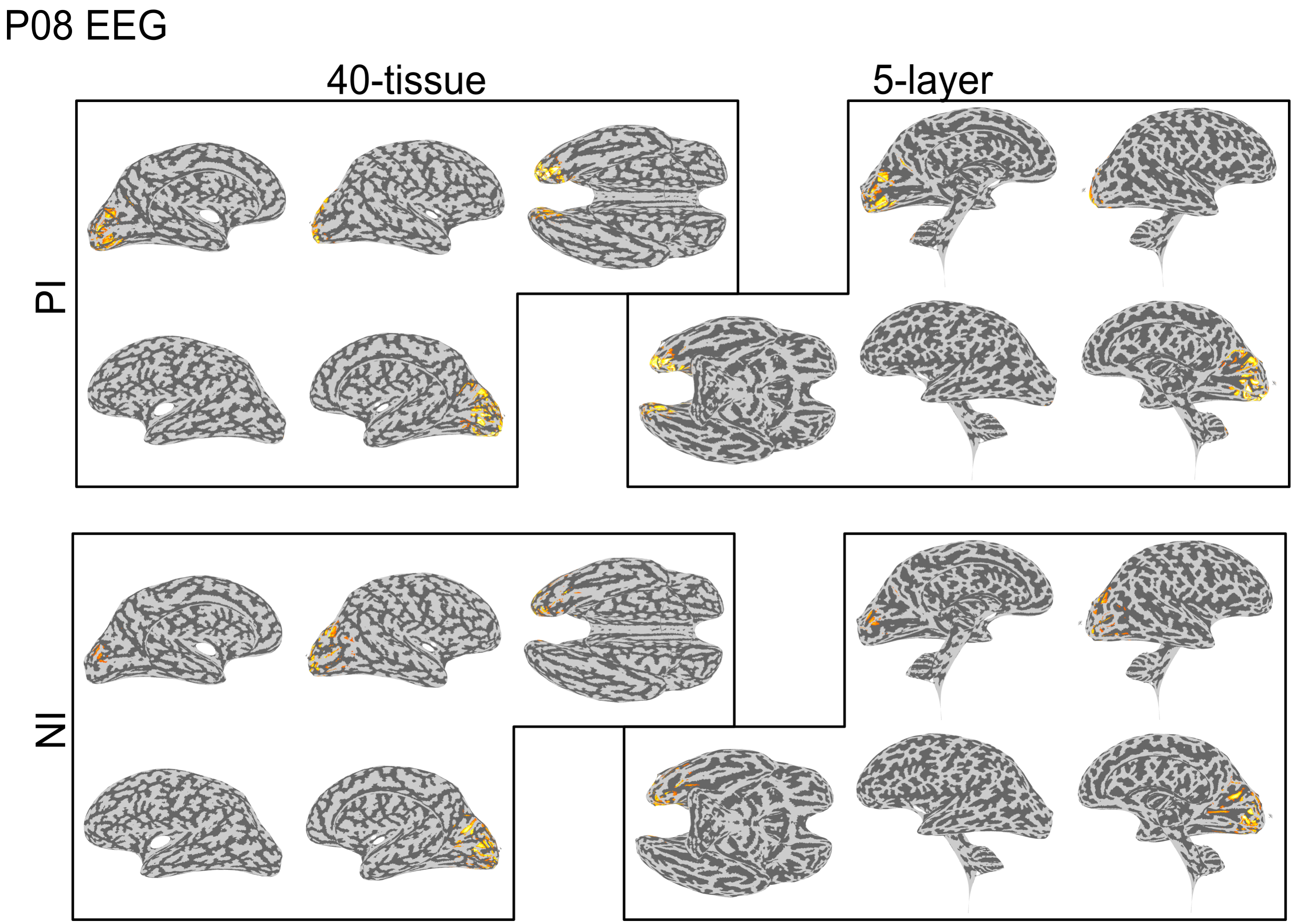


Figure S2.11: The EEG source localization solutions for subject P08 at each of the signal peaks ($P_I$ and $N_I$).

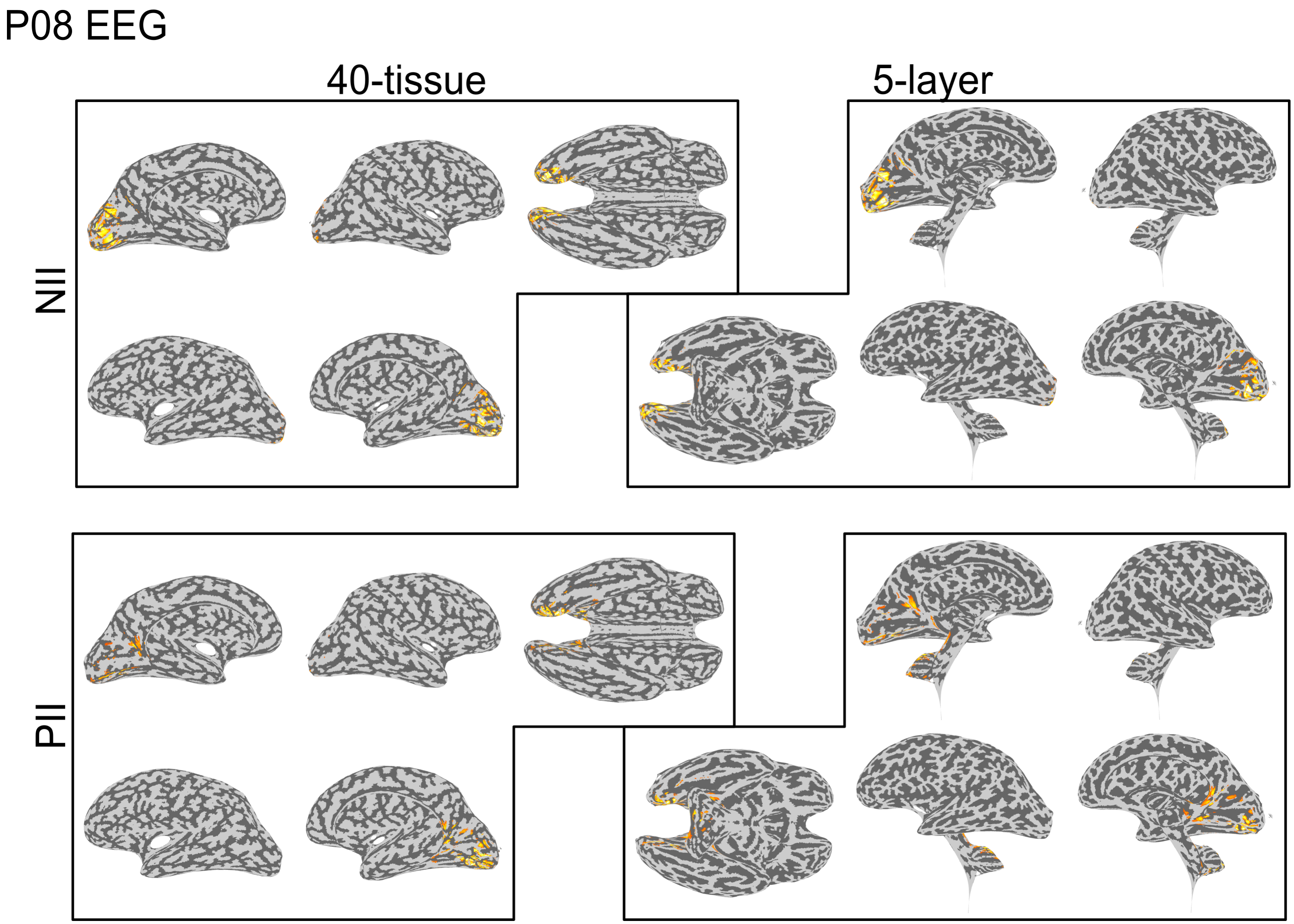


Figure S2.12: The EEG source localization solutions for subject P08 at each of the signal peaks ($N_{II}$ and $P_{II}$).

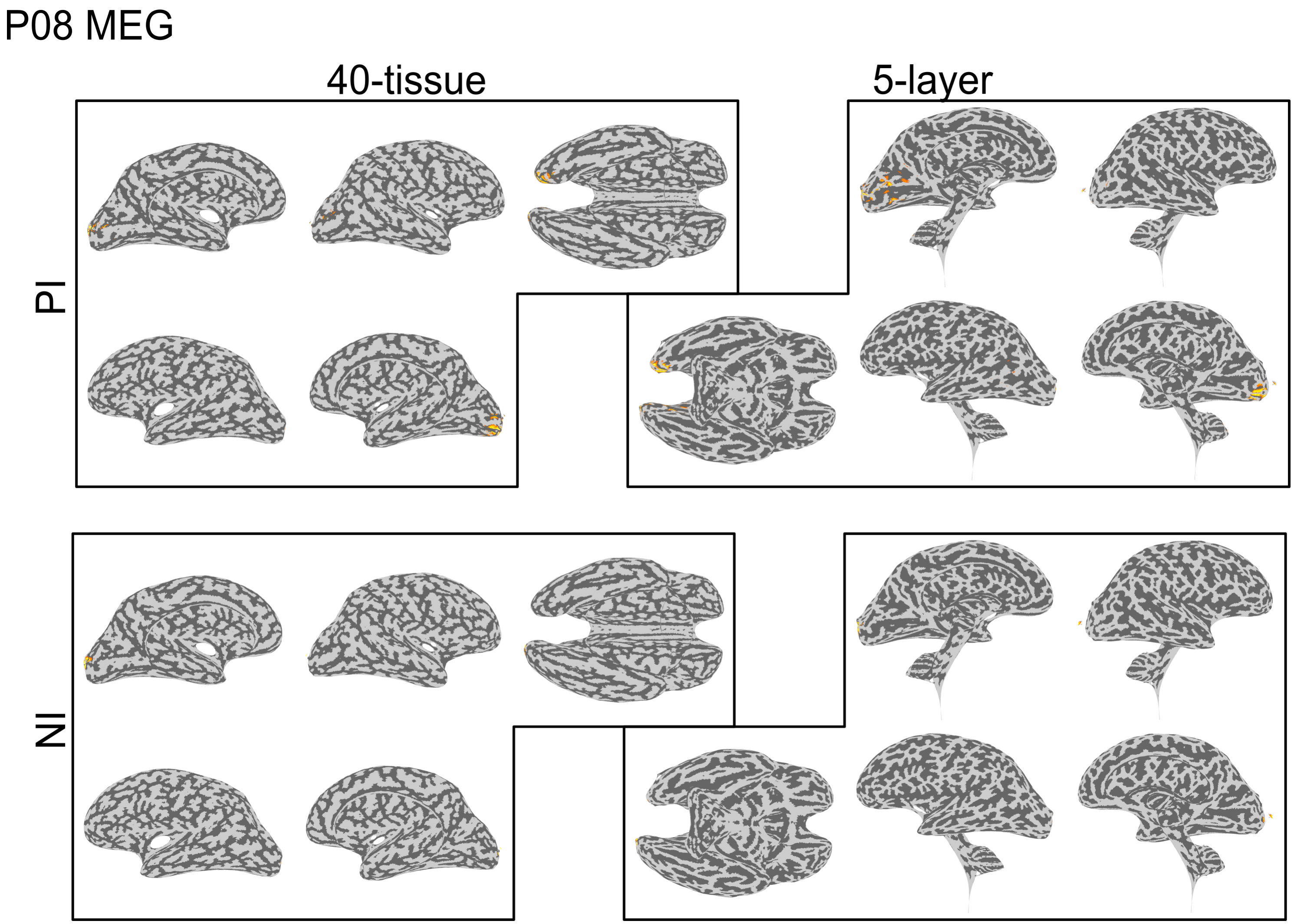


Figure S2.13: The MEG source localization solutions for subject P08 at each of the signal peaks ($P_I$ and $N_I$).

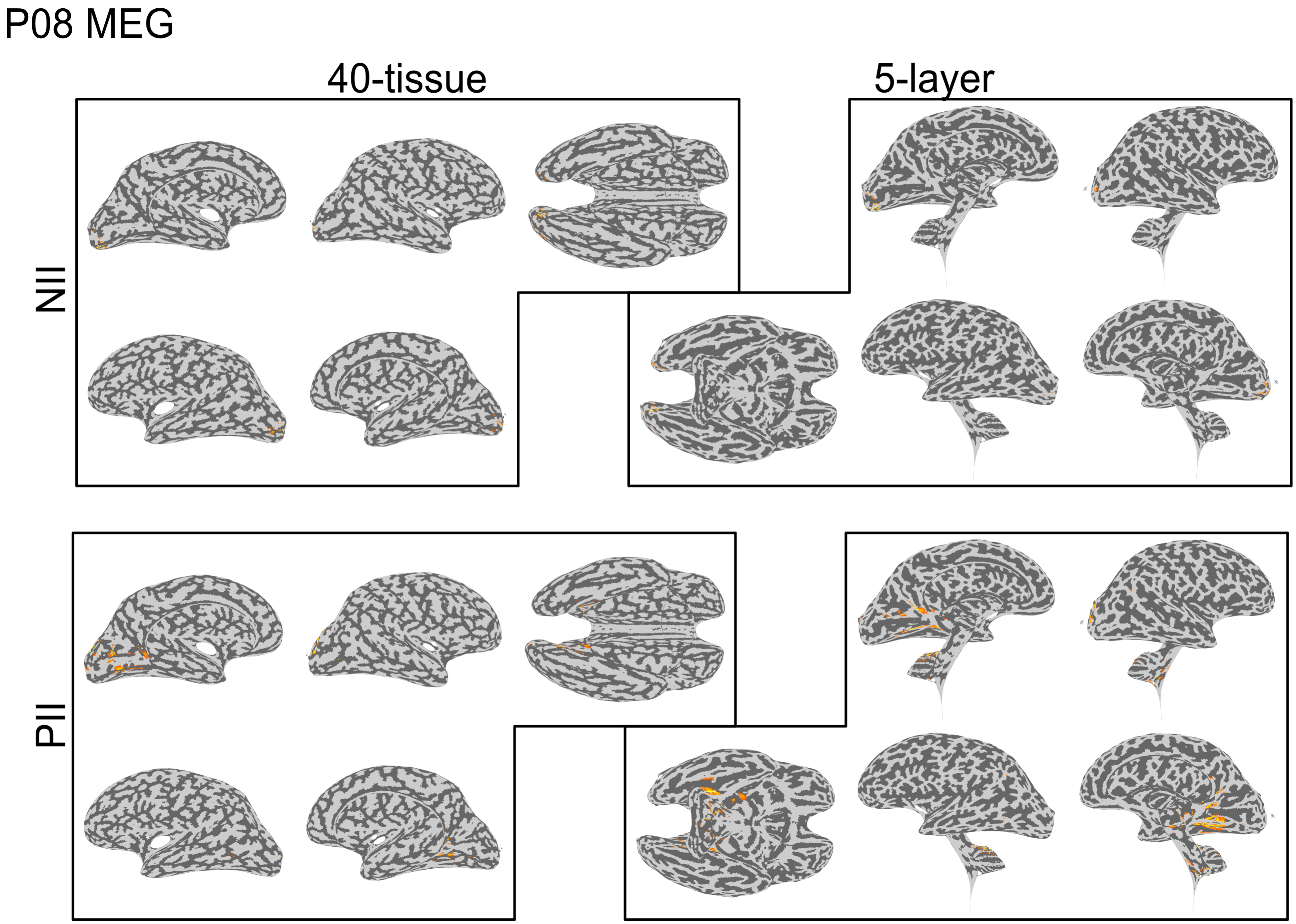


Figure S2.14: The MEG source localization solutions for subject P08 at each of the signal peaks ($N_{II}$ and $P_{II}$).

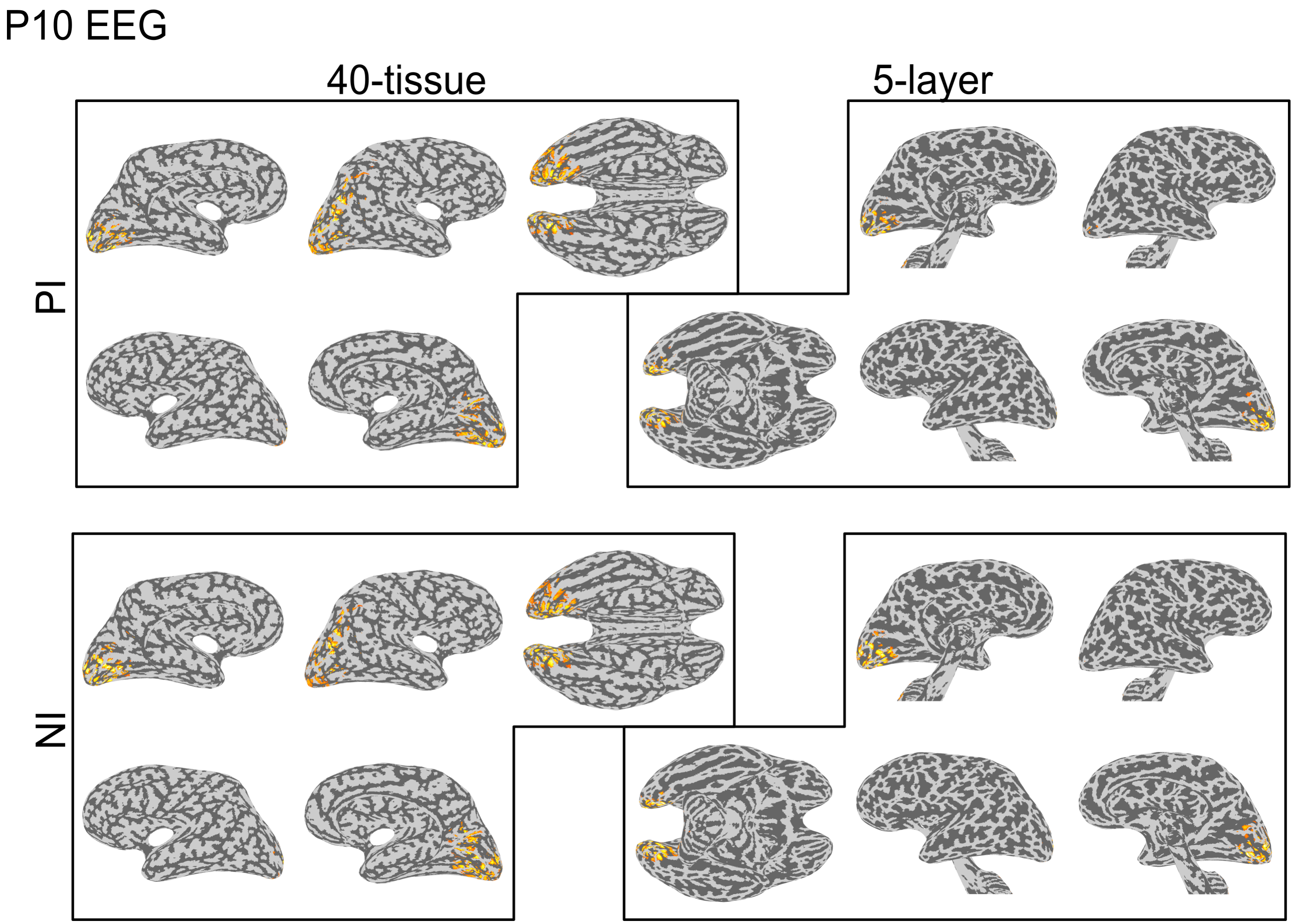


Figure S2.15: The EEG source localization solutions for subject P10 at each of the signal peaks ($P_I$ and $N_I$).

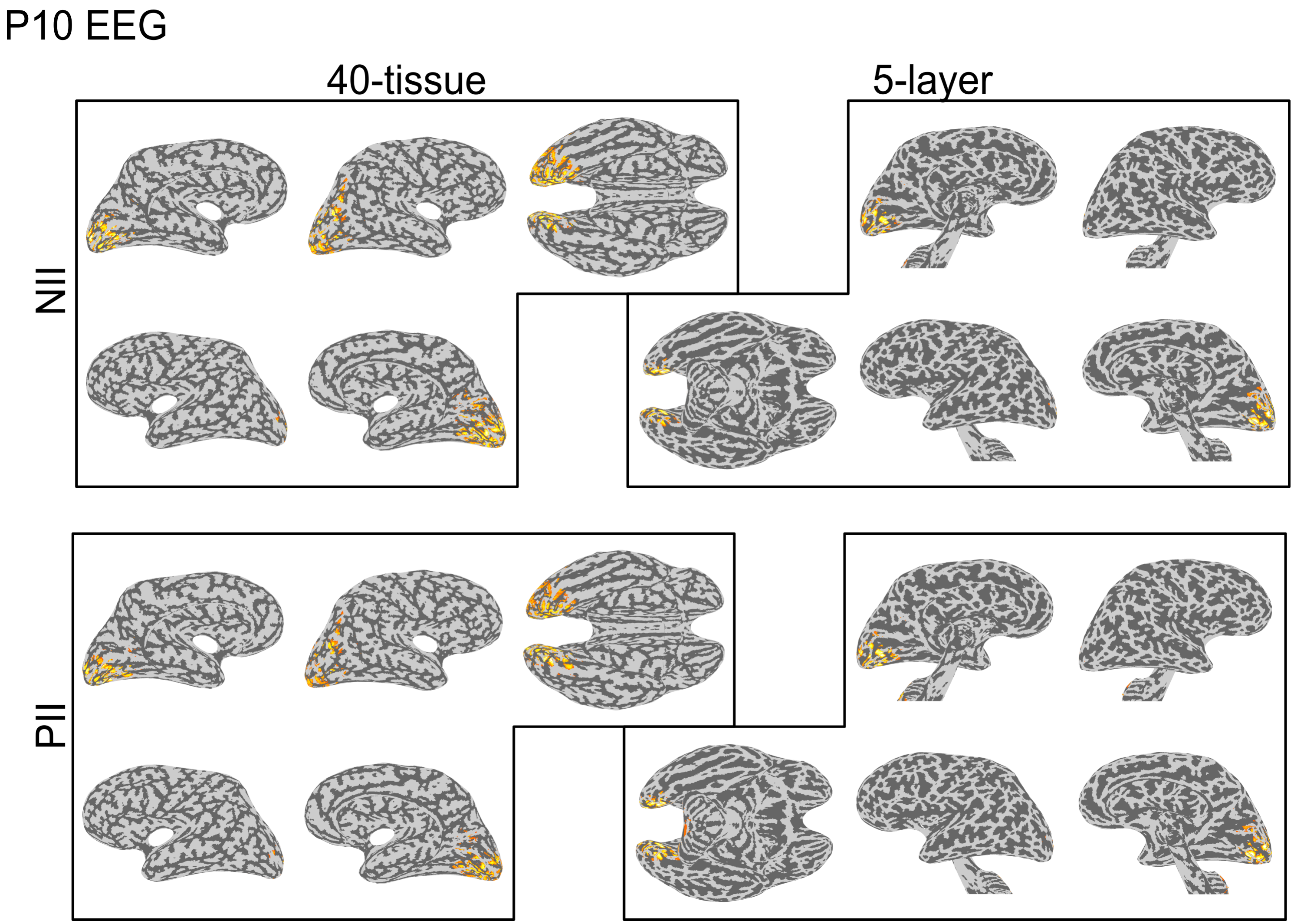


Figure S2.16: The EEG source localization solutions for subject P10 at each of the signal peaks ($N_{II}$ and $P_{II}$).

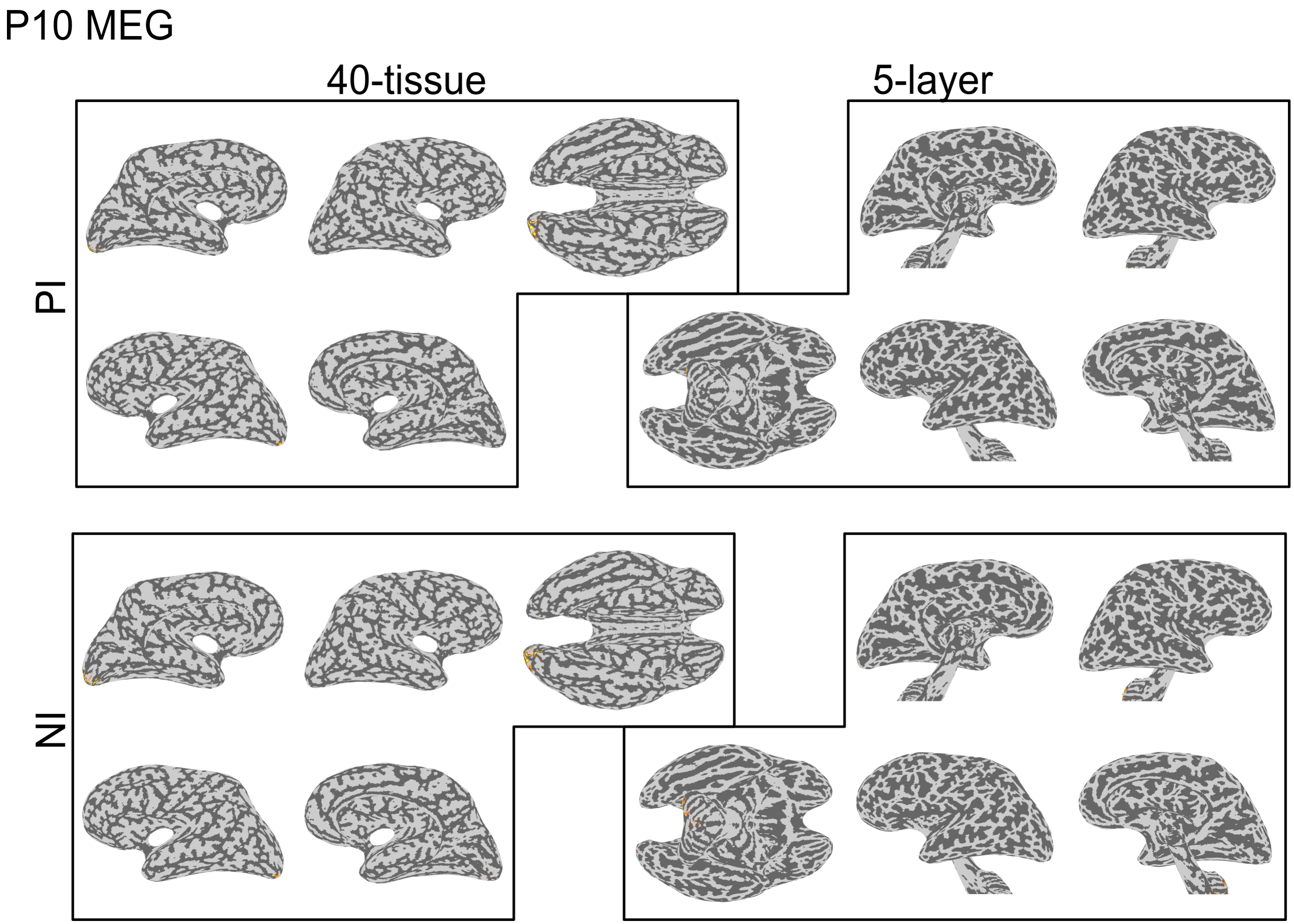


Figure S2.17: The MEG source localization solutions for subject P10 at each of the signal peaks ($P_I$ and $N_I$).

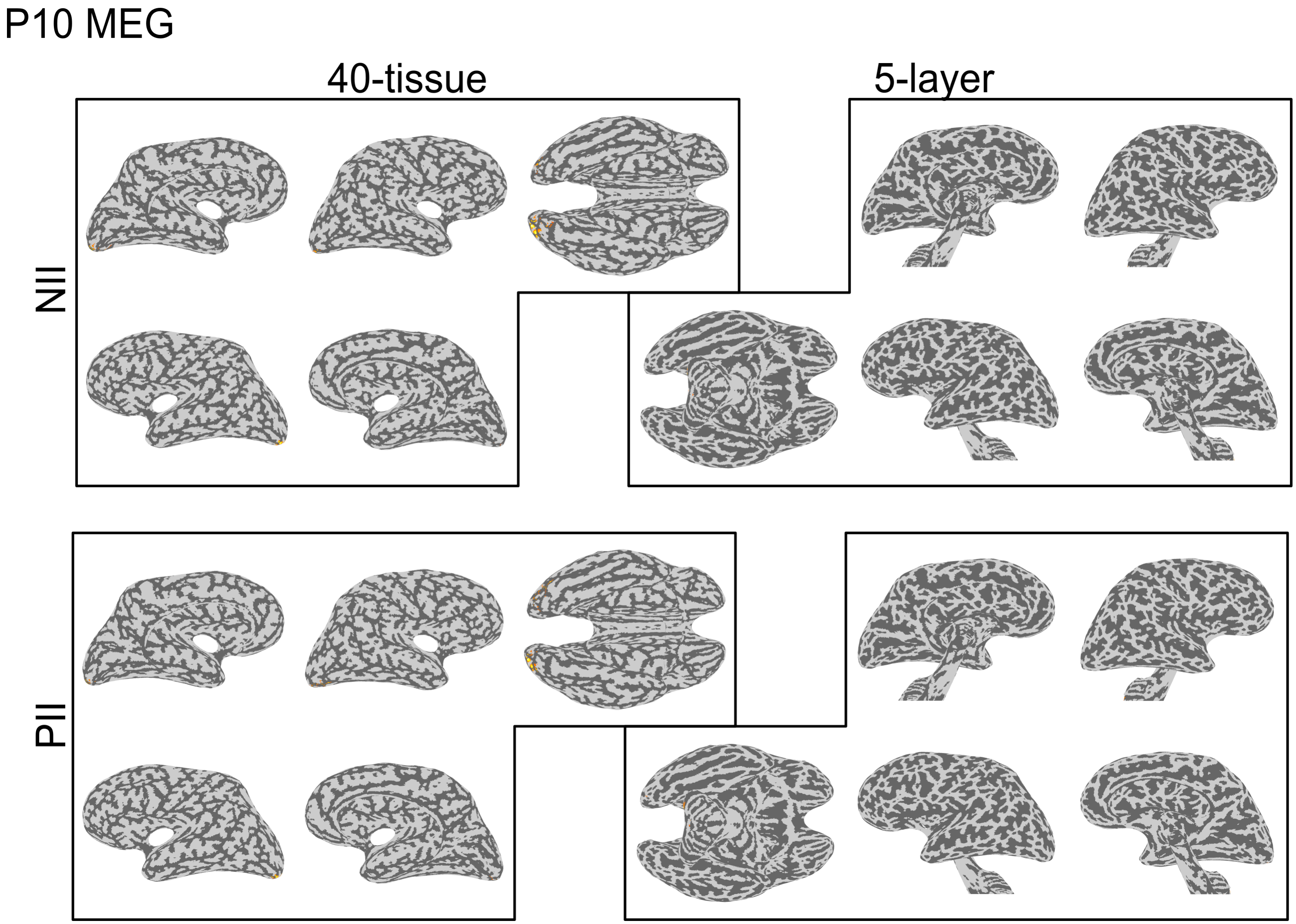


Figure S2.18: The MEG source localization solutions for subject P10 at each of the signal peaks ($N_{II}$ and $P_{II}$).

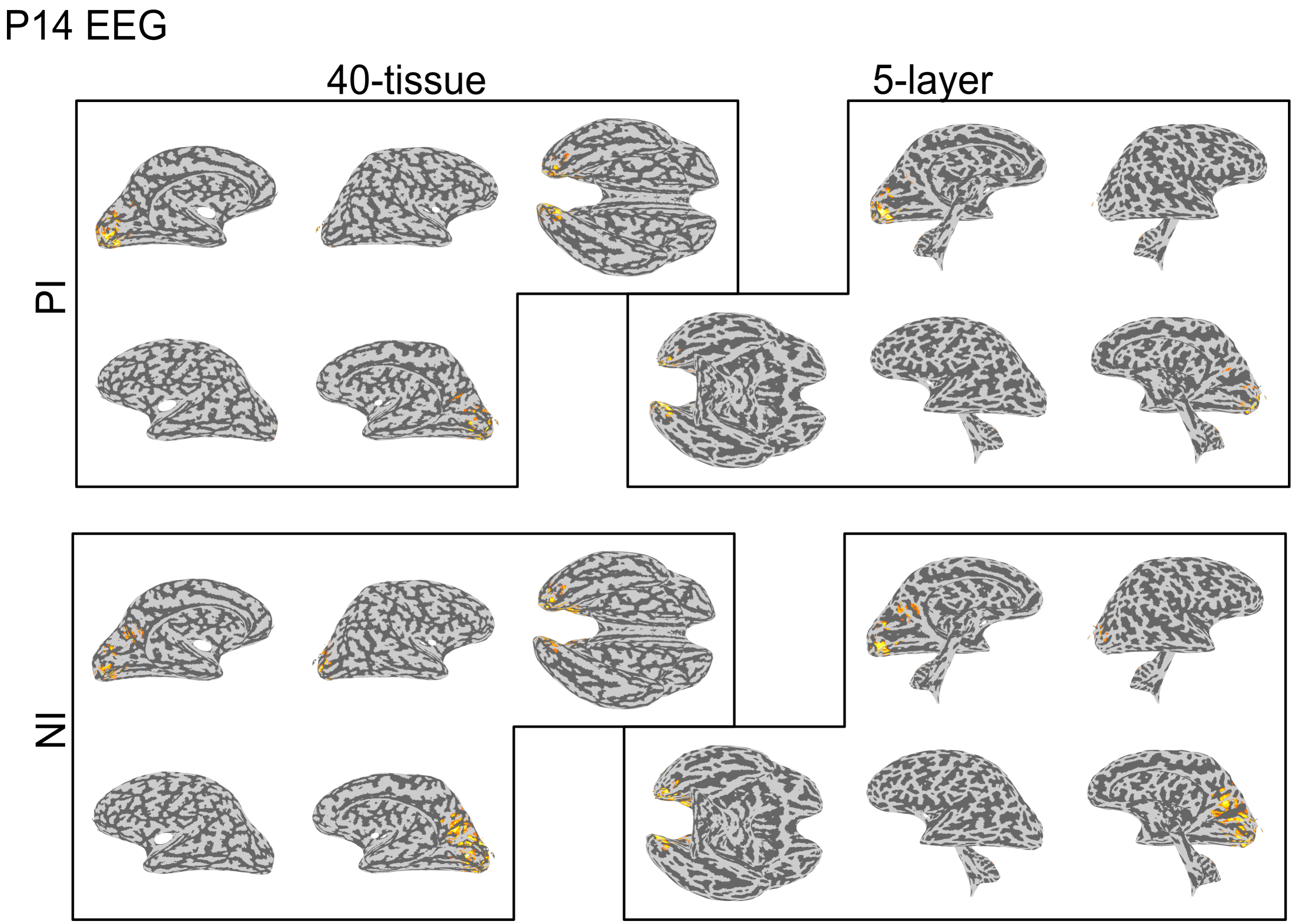


Figure S2.19: The EEG source localization solutions for subject P14 at each of the signal peaks ($P_I$ and $N_I$).

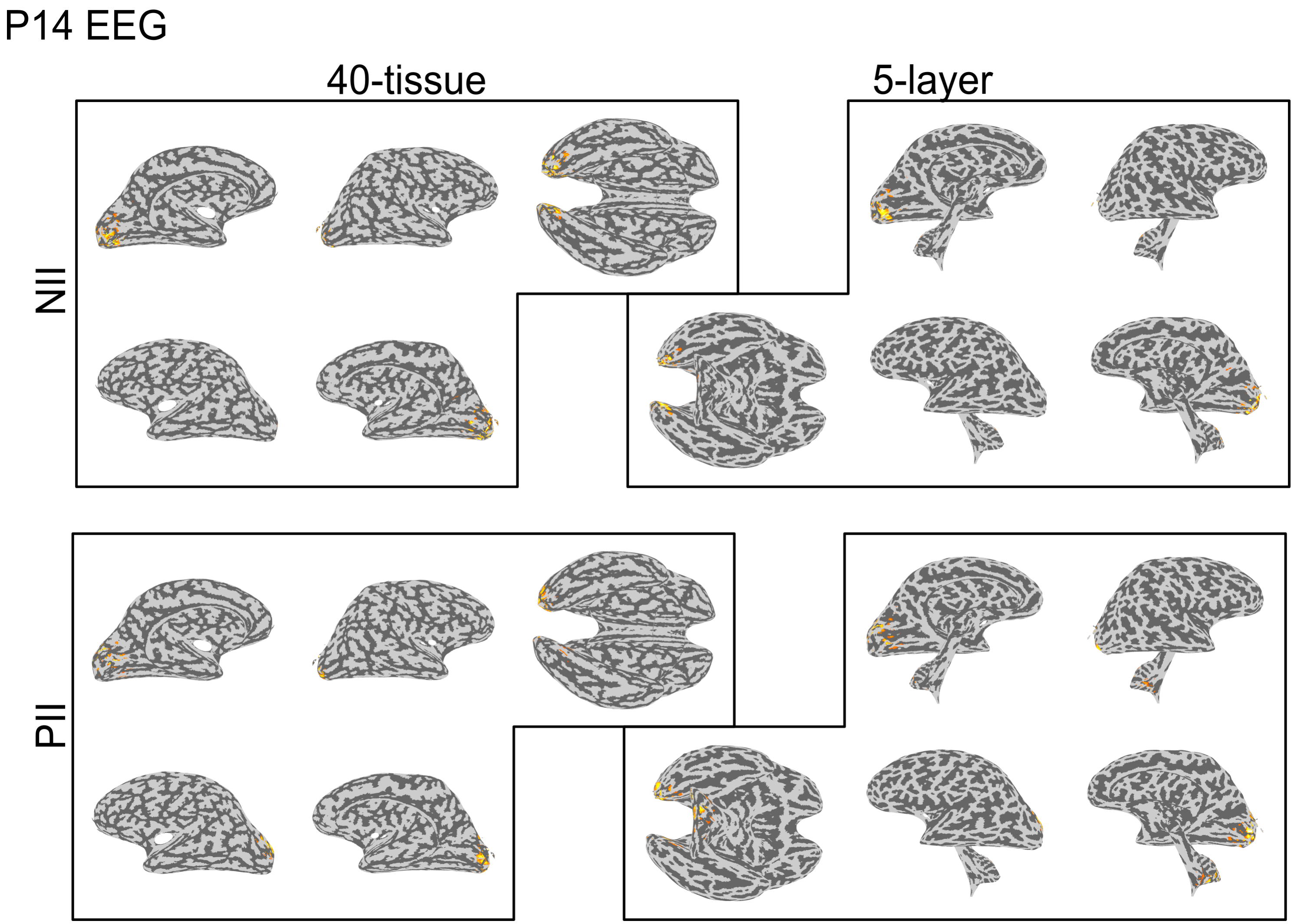


Figure S2.20: The EEG source localization solutions for subject P14 at each of the signal peaks ($N_{II}$ and $P_{II}$).

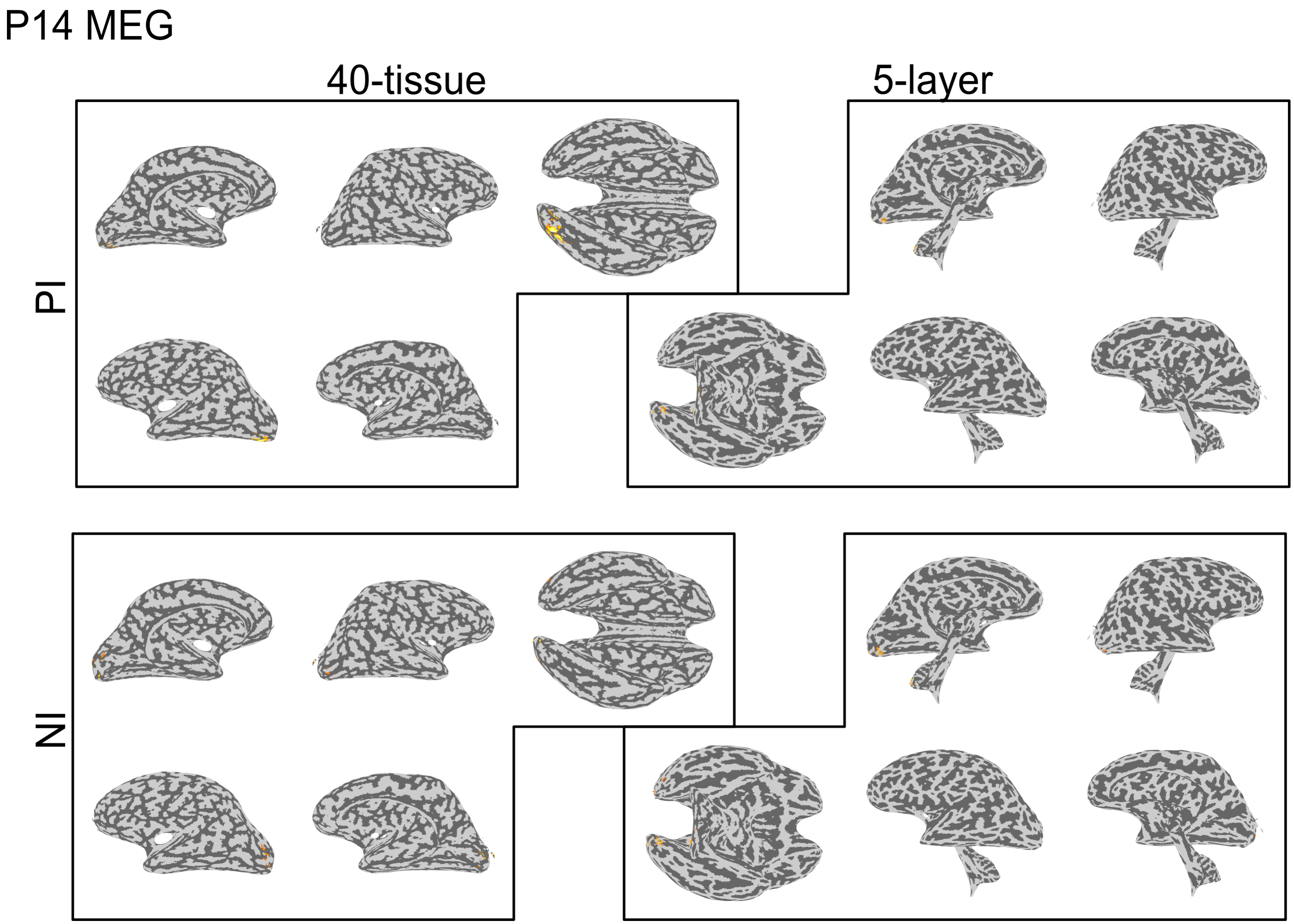


Figure S2.21: The MEG source localization solutions for subject P14 at each of the signal peaks ($P_I$ and $N_I$).

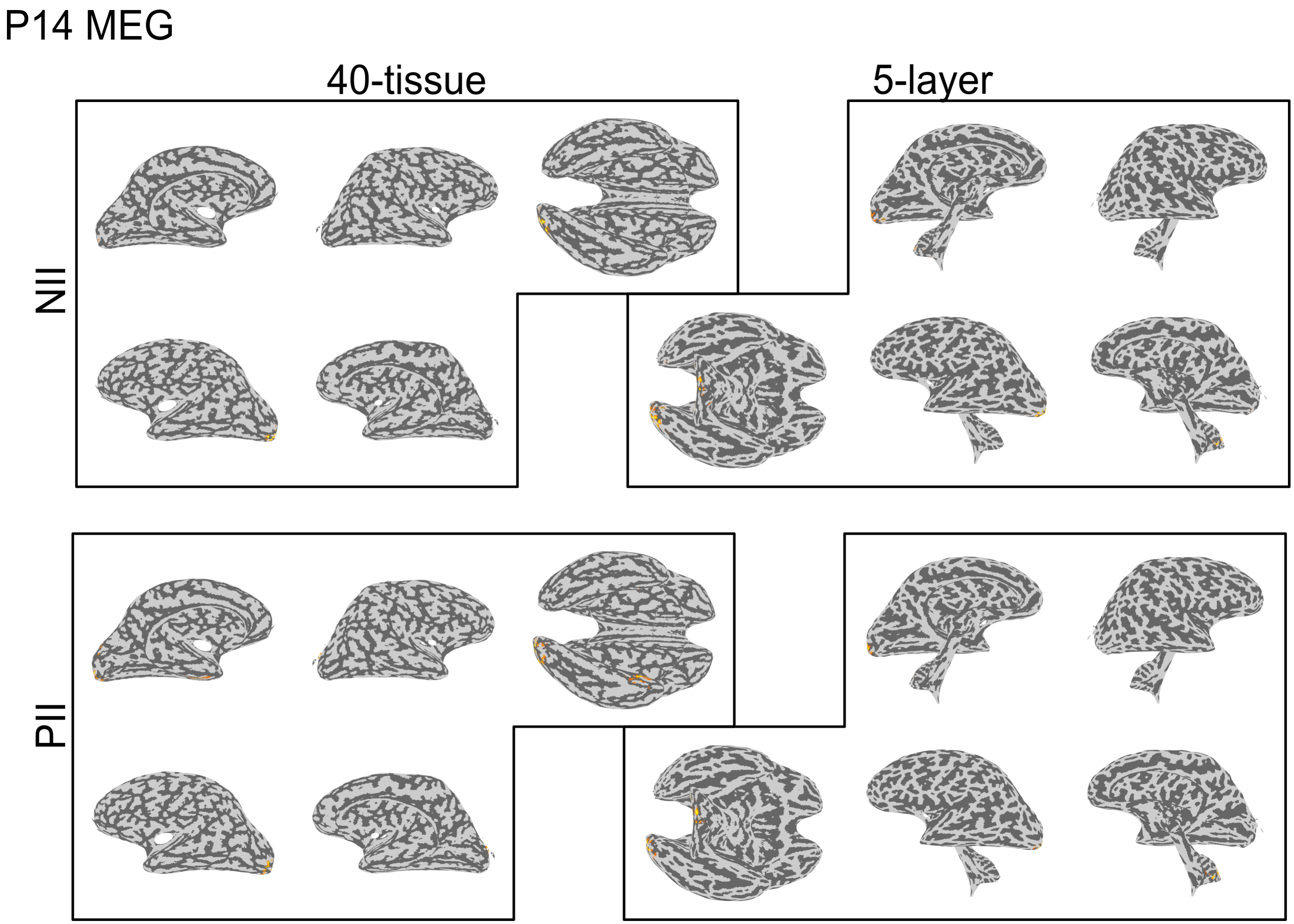


Figure S2.22: The MEG source localization solutions for subject P14 at each of the signal peaks ($N_{II}$ and $P_{II}$).

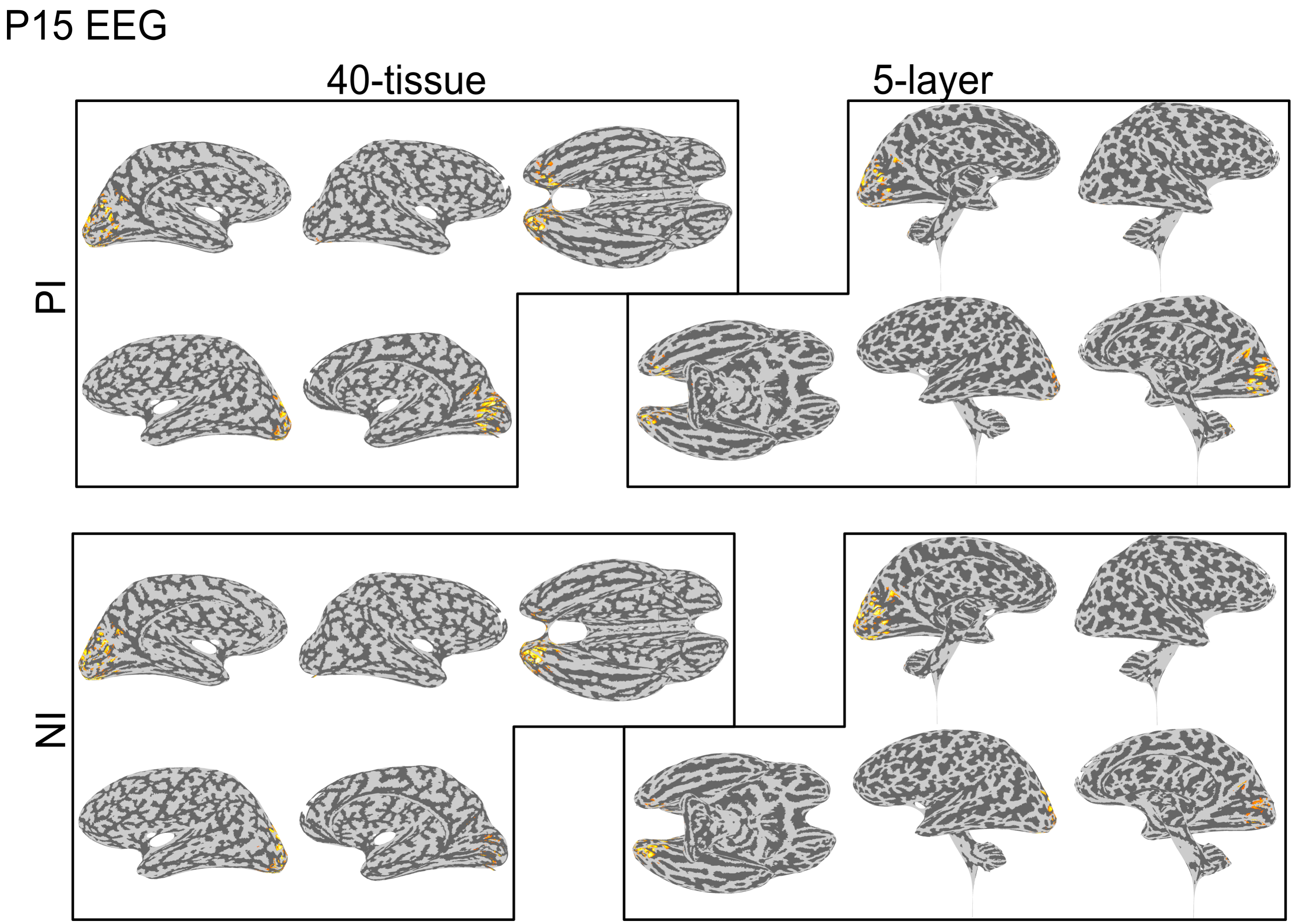


Figure S2.23: The EEG source localization solutions for subject P15 at each of the signal peaks ($P_I$ and $N_I$).

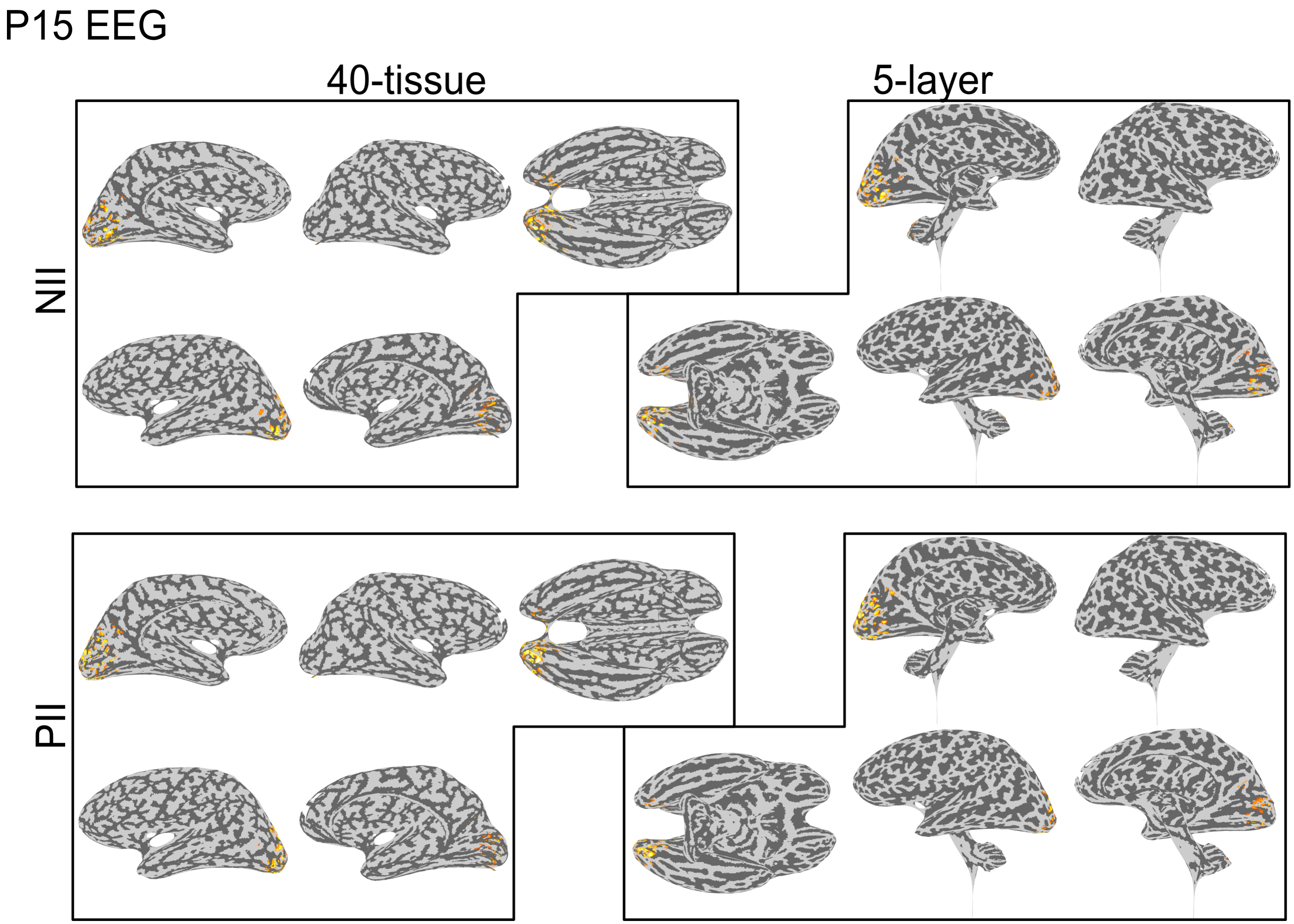


Figure S2.24: The EEG source localization solutions for subject P15 at each of the signal peaks ($N_{II}$ and $P_{II}$).

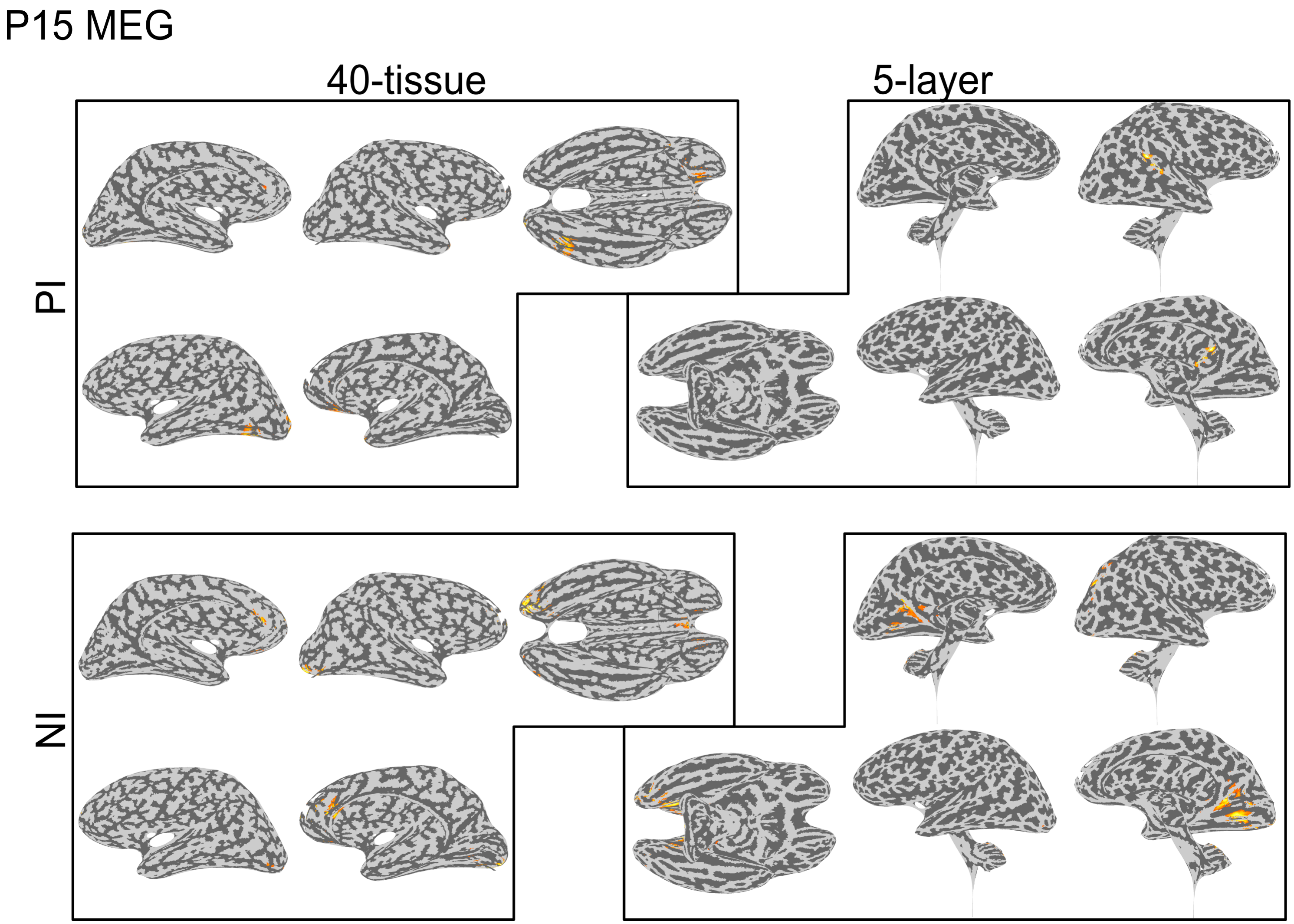


Figure S2.25: The MEG source localization solutions for subject P15 at each of the signal peaks ($P_I$ and $N_I$).

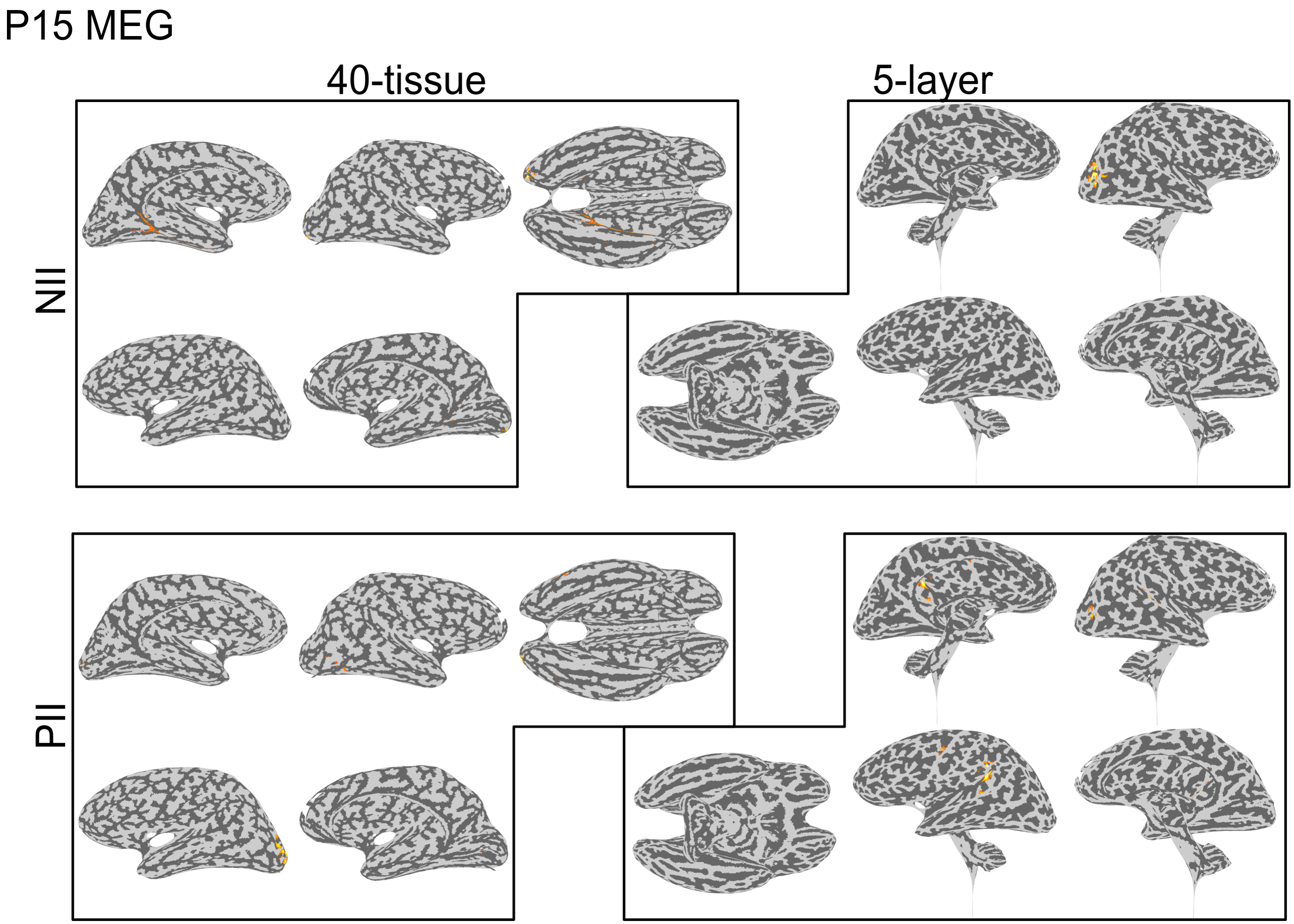


Figure S2.26: The MEG source localization solutions for subject P15 at each of the signal peaks ($N_{II}$ and $P_{II}$).
